# A lipid-laden macrophage niche drives immunosuppression in primary central nervous system lymphoma

**DOI:** 10.64898/2026.02.19.705289

**Authors:** Liang Hong, Subhashini Joshi, Min Liu, Shruti Sridhar, María Victoria Revuelta, Zi Yan Charmaine Ong, Sheng Cong Norbert Tay, Wei Xian Clemence Lai, Chartsiam Tipgomut, Patrick Jaynes, Yanfen Peng, Char Loo Tan, Susan Swee-Shan Hue, Siok-Bian Ng, Sanjay De Mel, Limei Poon, Yogeshini Batumalai, Jayalakshmi, Jennifer Brooks, Florian Hamberger, Brian J. Lane, Daniel Jimenez-Sanchez, Oliver Braubach, Meng Xiao, Jinmiao Chen, Qiang Pan-Hammarstrom, Ranga Sudharshan, Ashley P. Tsang, Arvind Rao, Evan T. Keller, Zachary Hawula, Melinda Burgess, Nella Tuczko, Colm Keane, Federico Ruiz Moreno, Nahuel Zamponi, Gero Knittel, H. Christian Reinhardt, Maurilio Ponzoni, Claudio Tripodo, Leandro Cerchietti, Anand Devaprasath Jeyasekharan

## Abstract

Primary central nervous system lymphoma (PCNSL) is histologically a subtype of diffuse large B-cell lymphoma (DLBCL), sharing genetic and transcriptomic similarity, but with distinct clinical features, particularly its confinement to the CNS and higher relapse risk. The microenvironmental basis for its divergence from systemic DLBCL remains unclear. Using spatial transcriptomic approaches (Xenium/GeoMx digital spatial profiling) in a comparative study of PCNSL (n=17) and DLBCL (n=76), we found that PCNSL, unlike systemic DLBCL, is dominated by an immunosuppressive macrophage compartment enriched for cholesterol-metabolism programs. In independent cohorts of PCNSL profiled by single-cell RNA sequencing, we validated the presence of a recurrent population of lipid-laden macrophages (LLMs): TREM2/GPNMB-expressing, lipid-remodeled cells transcriptionally distinct from resident microglia and consistent with an infiltrating monocyte origin, not previously characterized in CNS lymphoma. LLMs formed immunosuppressive niches with regulatory T cells, and using Cellscape hyperplex proteomic imaging we demonstrate that LLM-Treg spatial interactions are associated with chemotherapy response. To test whether LLMs are lymphoma-driven and functionally important, we developed an immunocompetent syngeneic PCNSL mouse model, driven by Myd88^L252P^ and Cd79b mutations with Bcl2 overexpression. Monocyte-derived macrophages in lymphoma-bearing brain regions acquired an LLM-like state, not seen in lymphoma-free brain regions or in splenic tumors driven by the same oncogenic lesions. TREM2-SYK signaling sustained this state, and SYK inhibition reversed its tumor-supportive activity *ex vivo*. These findings identify the LLM program as a targetable immunosuppressive myeloid state in CNS lymphoma.

**Key Points:**

1. PCNSL is enriched for TREM2^+^ lipid-laden macrophages (LLMs) forming immunosuppressive niches enriched relative to systemic DLBCL.
2. The first immunocompetent MCD-like-genotype PCNSL model recapitulates human LLMs; SYK inhibition reverses their tumor-supportive activity.

## Introduction

Primary central nervous system lymphoma (PCNSL) is a rare, highly aggressive subtype of diffuse large B-cell lymphoma (DLBCL) that arises within the brain, spinal cord, meninges, or eyes and rarely disseminates outside the CNS.^1,2^ Although histologically indistinguishable from systemic DLBCL, PCNSL is clinically and biologically distinct, with poor outcomes and no universally accepted standard of care in the relapsed or refractory setting.^2,3^ By cell of origin, PCNSL almost uniformly resembles the activated B-cell-like (ABC) subtype, which carries an inferior prognosis in systemic disease. At the genetic level, it correspondingly aligns with the MCD subtype, a predominantly ABC-associated genetic class defined by recurrent MYD88 and CD79B mutations that converge on NF-κB and B-cell receptor signaling.^4–9^ These alterations are shared with systemic ABC-DLBCL,^10–13^ and do not by themselves account for the strict CNS confinement of PCNSL, indicating that tumor-intrinsic lesions alone cannot explain its distinct behavior.

One explanation is that PCNSL is shaped by a specialized tumor microenvironment (TME). The PCNSL TME is immunosuppressive^14–17^ and develops within the unique CNS immune context, where surveillance is tightly regulated through neuroimmune crosstalk.^18,19^ Although malignant B-cell programs and T-cell dysfunction in PCNSL are increasingly defined,^15–17,20^ the abundant tumor-associated macrophage compartment remains comparatively understudied, despite accounting for approximately 10%-13% of captured cells in PCNSL tissues.^20–22^ Prior studies have relied on a small number of canonical lineage and polarization markers assessed within a binary M1/M2 framework,^23–25^ yielding conflicting prognostic associations and little insight into macrophage spatial organization, tumor-proximal identity, or cell state.^26–32^ This gap is important because tissue-resident microglia and infiltrating monocyte-derived macrophages (MDMs) are distinct regulators of CNS disease,^33^ and in glioma occupy separate niches linked to survival.^34,35^ Spatial studies have also linked macrophage programs to clinical outcomes in DLBCL and PCNSL.^36,37^ A macrophage-focused, spatially resolved comparison of PCNSL with systemic DLBCL is therefore needed.

Determining whether PCNSL macrophage states are lymphoma-driven and functionally important requires a genetically faithful, immunocompetent mouse model. Existing immunocompetent mouse models of CNS lymphoma rely on transplantable lines such as A20 and BAL17, which lack the hallmark genetic lesions that define human PCNSL,^38,39^ limiting their ability to model disease-specific, genotype-driven microenvironmental biology.

Here, we integrated spatial transcriptomics, spatial proteomics, and single-cell profiling to define how the tumor-associated macrophage compartment differs between PCNSL and systemic DLBCL. This revealed a lipid-laden macrophage (LLM) state characteristic of PCNSL that forms immunosuppressive niches with regulatory T cells, with the spatial enrichment of Tregs around LLMs tracking with chemotherapy response. To move from association to causation, we then developed the first immunocompetent, MCD-like-genotype-driven syngeneic model of PCNSL and used it to show that LLMs are induced specifically in lymphoma-bearing brain, but not in genotype-matched systemic tumors, establishing the CNS site as a key variable, with sensitivity to SYK inhibition. Together, these human and experimental data nominate LLMs as a CNS-restricted, lymphoma-driven macrophage state and a candidate therapeutic target.

## Methods

Full methods, including all computational parameters, are provided in supplemental Methods.

### Patient samples and study design

This study was approved by the Singapore NHG Domain Specific Review Board study protocol 2015/00176, and was conducted in accordance with relevant ethical regulations. Pretreatment formalin-fixed, paraffin-embedded (FFPE) biopsies were obtained from 24 patients with primary central nervous system lymphoma (PCNSL) and 76 systemic diffuse large B-cell lymphoma (DLBCL). Tonsil controls were obtained from the NUH cohort under the same waiver. Diagnoses were confirmed by histopathologic review. Tissue for the University of Queensland single-cell cohort was obtained from the Plymouth NHS Trust through BRAIN UK.^40^ Assays used platform-specific subsets of available tissue because of differences in residual tissue, section quality, tissue loss during processing, and sectioning requirements. Patient overlap and platform assignment are provided in Supplemental Table 1.

### Spatial transcriptomics and proteomics

Tissue microarrays were profiled by Xenium In Situ (10x Genomics; 380-gene Immuno-Oncology panel; 17 PCNSL and 76 DLBCL cases) and by GeoMx DSP whole-transcriptome profiling (17 PCNSL, 64 DLBCL) using CD3, CD20, and CD68 masks to capture compartment-specific areas of interest (AOIs). GeoMx data were processed with standR, limma, and vissE, with BayesPrism deconvolution against the MoMac-VERSE atlas.^41^

Spatial proteomics used CellScape (35-marker panel including lipid-associated macrophage markers: GPNMB, TREM2, CD36)^42^ on 16 PCNSL and 5 tonsil samples. GPNMB defines the GBM-derived LLM signature^43^ and gave the most consistent staining, so we used GPNMB^+^CD68^+^ cells to identify LLM-like macrophages. Cells were segmented in Cellpose and analyzed in QuPath and R; LLM abundance, pair-correlation, and treatment-response comparisons were performed at the patient level.

### Single-cell RNA-seq

Four scRNA-seq cohorts were analyzed: two published discovery datasets, PCNSL (Liu et al, n = 8)^15^ and DLBCL (Ye et al, n = 17)^44^; validation cohort 1, an in-house University of Queensland cohort (13 PCNSL, 4 DLBCL); and validation cohort 2, published PCNSL and reactive lymph node samples (Kobayashi et al, 7 PCNSL, 2 reactive lymph nodes).^45^ Data were processed in Seurat with quality control, batch correction, and myeloid/lymphoid subclustering. The LLM signature was derived through integration of published lipid-associated and glioblastoma (GBM) LLM programs.^43,46^ LIANA+ was used for ligand-receptor analysis.^47^

### Immunocompetent MCD-driven PCNSL model

Murine MCD DLBCL cells (MCD_P2; Myd88^L252P^, Cd79b-mutant, Bcl2-overexpressing) were implanted into the left hemisphere of wild-type male C57BL/6 mice; splenic implants provided a matched systemic comparator. On day 15, CNS tumor-site, contralateral non-tumor-site, and splenic tumor tissues were collected for single-cell RNA-seq (three mice pooled per condition), integrated in Seurat with Harmony. For functional assays, primary Cd11b^+^ brain myeloid cells were cultured in MCD_P2-conditioned medium ± the SYK inhibitor BAY 61-3606, then co-cultured with CellTrace Violet-labeled MCD_P2 cells; proliferation and Trem2/Cd163 expression were assessed by flow cytometry. All animal procedures were conducted in accordance with NIH guidelines and approved by the Institutional Animal Care and Use Committee (IACUC) of Weill Cornell Medicine (protocol #2015-0019).

### Statistical analysis

Analyses used R v4.4.3. Xenium comparisons were at the core/ROI level and GeoMx at the patient level (one AOI each patient); single-cell comparisons used patient-level summaries for inference where identifiers were available. Murine single-cell data were interpreted descriptively because cells were pooled per condition. Multiple comparisons were adjusted by Benjamini-Hochberg (FDR < 0.05).

## Results

### PCNSL exhibits an immunosuppressive, macrophage-enriched landscape

We compared PCNSL and systemic DLBCL immune landscapes in NUH tissue microarrays comprising 76 DLBCL patients (171 cores) and 17 PCNSL patients (17 cores), using the Xenium Immuno-Oncology panel and canonical marker-based phenotyping (Figure 1A-B; supplemental Figure 1A-B).

**Figure 1.**
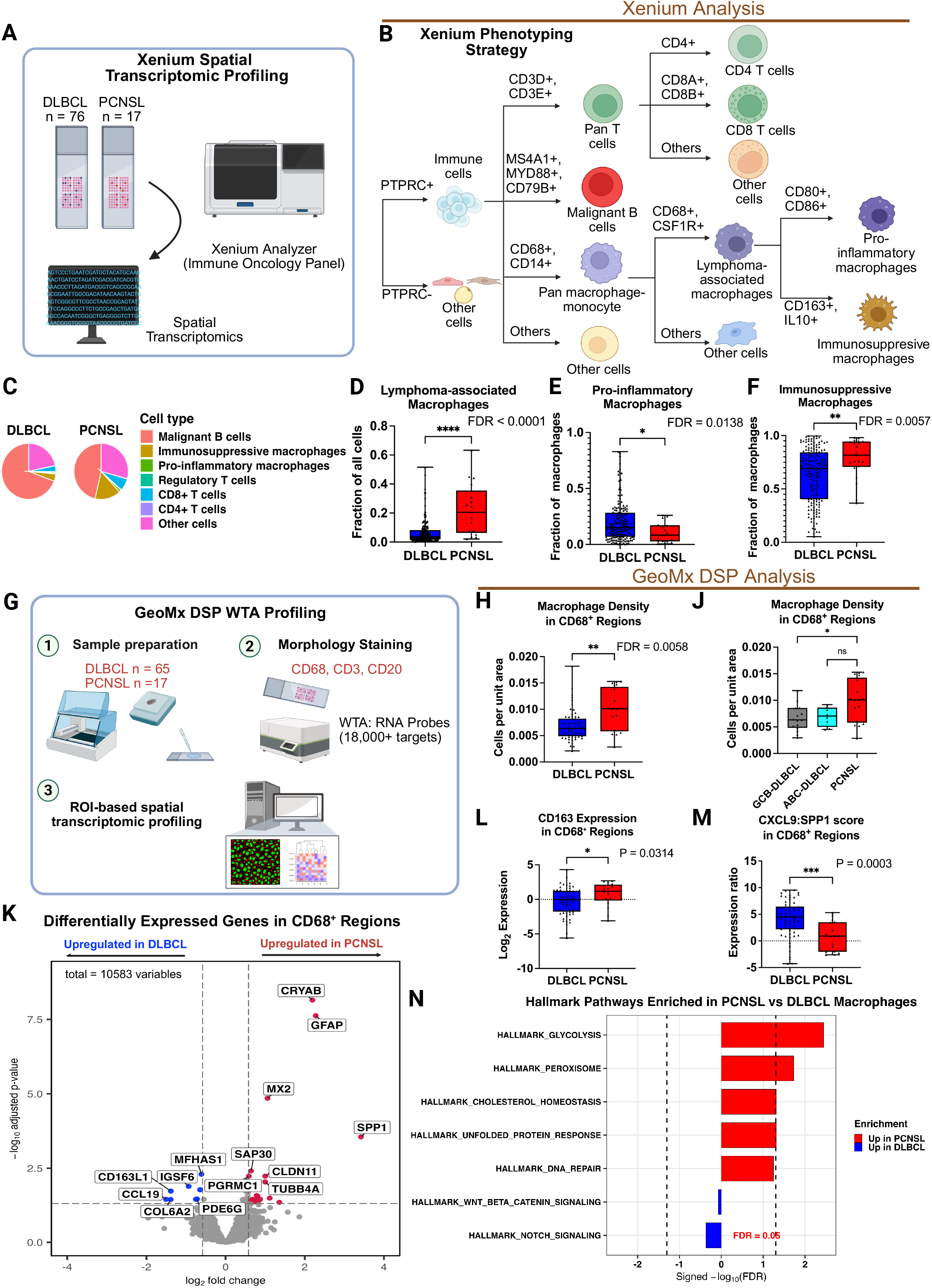
Spatial phenotyping reveals divergent immune composition and distinct macrophage transcriptional programs in PCNSL versus systemic DLBCL. (A) Overview of the Xenium spatial profiling workflow. (B) Marker-based phenotyping strategy for major immune lineages and for T-cell and macrophage/myeloid subsets. (C) Relative cell-type composition in PCNSL and DLBCL. (D) Macrophage proportions in PCNSL versus DLBCL. (E) Proinflammatory macrophage/myeloid proportions, defined as CD80^+^CD86^+^ cells. (F) Immunosuppressive macrophage/myeloid proportions, defined as CD163^+^IL10^+^ cells. In panels D-F, each point represents 1 TMA core. (G) Overview of the GeoMx Whole Transcriptome Atlas workflow. The initial cohort comprised 17 PCNSL and 64 DLBCL AOIs; after quality control, 17 PCNSL and 62 DLBCL AOIs were retained for downstream analysis. (H, J) Macrophage density in CD68^+^ AOIs, calculated as nuclei count divided by AOI area, comparing PCNSL with systemic DLBCL overall (H) and with DLBCL stratified by cell-of-origin subtype (J). (K) Differential expression in PCNSL versus DLBCL CD68⁺ AOIs; differentially expressed genes were defined as |log_2_ fold change| > 0.58 and adjusted P < 0.05. (L) CD163 expression in CD68^+^ AOIs, shown as log_2_-transformed TMM-normalized counts. (M) CXCL9:SPP1 expression ratio in CD68^+^ AOIs. (N) Hallmark gene set testing in CD68⁺ AOIs from PCNSL and DLBCL. Box-and-whisker plots show the median and interquartile range, with whiskers extending to minimum and maximum. Each point represents 1 AOI. Statistical analyses are described in the method section.

PCNSL showed lower B-cell content and higher macrophage abundance than DLBCL (median proportions: malignant B cells, 36.4% vs 72.6%; T cells, 12.9% vs 5.9%; macrophages, 20.5% vs 3.7%; Figure 1C-D; supplemental Figure 2A-C; Supplemental Table 2). In both entities, the B-cell compartment is predominantly neoplastic,^48^ and is hereafter referred to as malignant B cells. Although total T-cell abundance did not differ significantly (FDR = 0.0609), the CD4:CD8 ratio was lower in PCNSL (P = 0.0008; supplemental Figure 2D-E). Within the macrophage compartment, pro-inflammatory phenotypes were enriched in DLBCL (FDR = 0.0138), whereas immunosuppressive phenotypes were enriched in PCNSL (FDR = 0.0057; Figure 1E-F).

GeoMx DSP whole-transcriptome profiling retained CD68⁺ AOIs from 17 PCNSL and 62 DLBCL patients after quality control, confirming the expected compartment enrichment (Figure 1G; supplemental Figure 3A-H). Consistent with the Xenium findings, CD68^+^ region cell density was higher in PCNSL than in DLBCL overall (FDR = 0.0058). In subtype-stratified comparisons, this difference reached significance against GCB-DLBCL but not against ABC-DLBCL, although the direction was consistent (Figure 1H-J). CD3^+^ region density did not differ between groups (supplemental Figure 3J). Differential expression analysis of CD68^+^ AOIs identified 20 PCNSL-upregulated and 8 DLBCL-upregulated genes (|logFC| > 0.58, adjusted P < 0.05; Figure 1K; Supplemental Table 3). CD68⁺ regions in PCNSL showed increased SPP1 and CRYAB but decreased CD163L1, IGSF6, and CCL19, whose predominant source in CNS lymphoma was recently identified as pericytes.^49^

CD163 expression was higher in PCNSL CD68^+^ AOIs (P = 0.0314; Figure 1L). As the CXCL9:SPP1 ratio distinguishes CXCL9-associated inflammatory from SPP1-associated immunoregulatory macrophage programs,^50^ we examined this ratio and found it significantly lower in PCNSL CD68⁺ AOIs (P = 0.0003; Figure 1M), consistent with an immunoregulatory skew. Gene set enrichment analysis further showed enrichment for glycolysis and lipid-metabolic programs, notably cholesterol homeostasis and peroxisome pathways (Figure 1N; supplemental Figure 3K), a lipid-loading signature characteristic of the immunosuppressive macrophages described in other tumors.^43,51^

### Macrophages in PCNSL have a TREM2/LLM-like immunosuppressive imprint

To resolve macrophage functional subtype differences between PCNSL and DLBCL, we projected the CD68^+^ differentially expressed genes onto the MoMac-VERSE monocyte and macrophage atlas^41^ (Figure 2A; supplemental Figure 4A). PCNSL-upregulated genes mapped to ISG monocytes, IL1B monocytes, and TREM2 macrophages, whereas DLBCL-upregulated genes mapped to dendritic cells, CD16^+^ monocytes, and IL4I1 macrophages (Figure 2B-D; supplemental Figure 4B-C).

**Figure 2.**
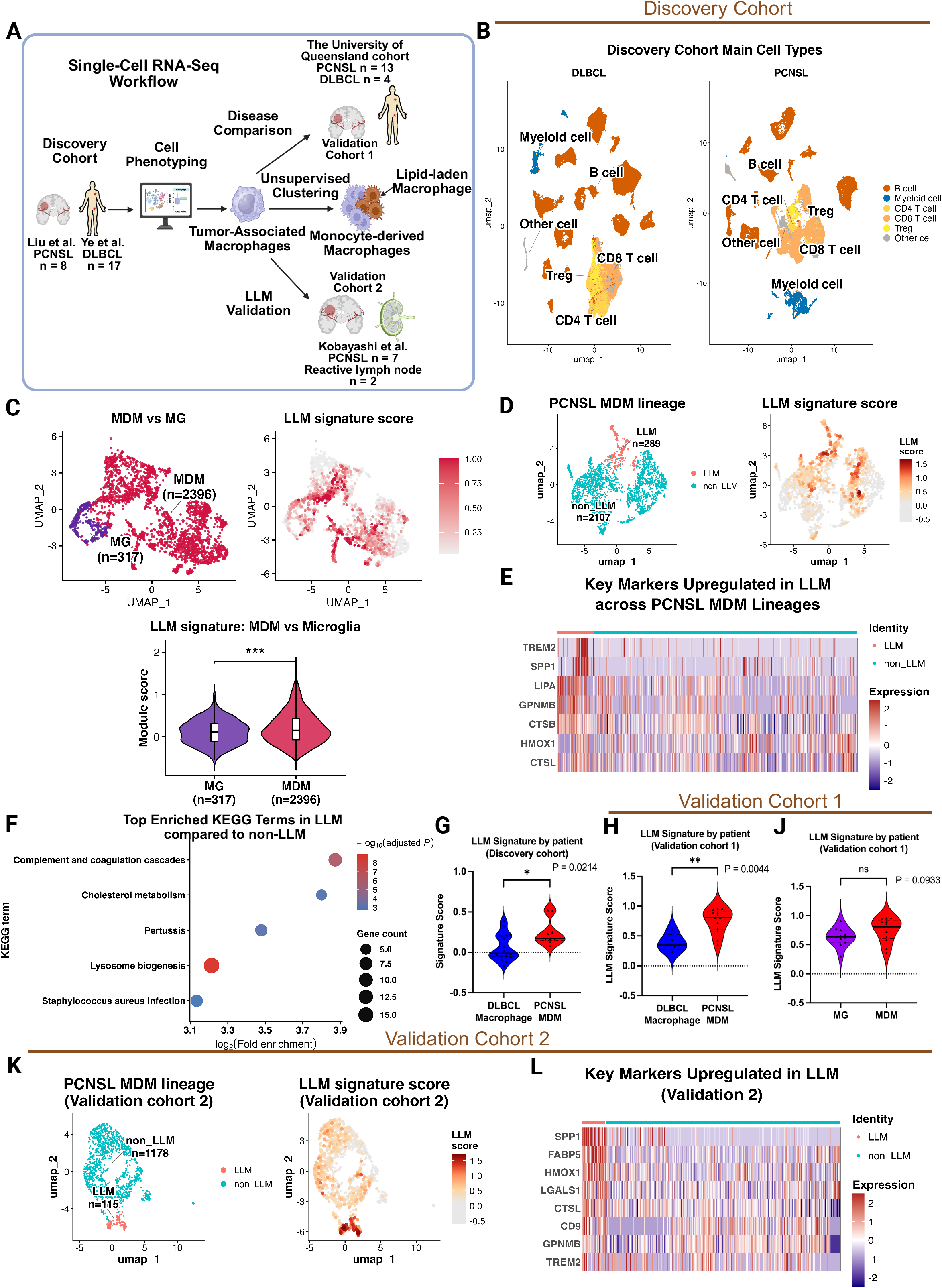
Signature projection and deconvolution of CD68^+^ AOIs reveal distinct macrophage states in PCNSL versus systemic DLBCL. (A) Overview of the MoMac-VERSE signature-projection and BayesPrism deconvolution workflow. (B) Difference in macrophage signature scores between PCNSL and DLBCL across MoMac-VERSE subtypes, calculated as PCNSL minus DLBCL. Points indicate median delta scores and horizontal bars indicate 95% bootstrap confidence intervals; positive values indicate relative enrichment of the PCNSL signature. Bootstrap intervals reflect within-subtype cell-level variability and do not constitute patient-level inference or hypothesis testing. (C-D) Projection of genes upregulated in DLBCL (C) or PCNSL (D) CD68**^+^** AOIs onto the MoMac-VERSE reference atlas, highlighting the top-ranked subsets for each signature. (E) BayesPrism-estimated macrophage-state proportions in PCNSL (n = 17 AOIs) and DLBCL (n = 62 AOIs). Only states with a median estimated abundance greater than 1% across all AOIs are shown. (F) Correlation between TREM2-macrophage and lipid-associated macrophage module scores in PCNSL CD68**^+^** AOIs. Module scores were calculated as the mean of gene-wise z scores. The line indicates the least-squares fit, shown for visualization. (G) Provenance of the macrophage lipid gene signatures used in this study. The lipid-associated macrophage module was derived from adipose tissue LAMs (Jaitin et al., Cell 2019)^46^; the lipid-laden macrophage (LLM) signature was derived from GPNMB^high^ macrophages in glioblastoma (Kloosterman et al., Cell 2024)^43^. Full gene lists are in Supplemental Table 4. (H) Lipid-associated module scores in PCNSL versus DLBCL CD68**^+^** AOIs. (J) GBM-derived GPNMB^high^ LLM signature scores in PCNSL versus DLBCL CD68**^+^** AOIs. (K) GBM-derived GPNMB^high^ LLM signature scores in PCNSL versus DLBCL CD68**^+^** AOIs, with DLBCL stratified by cell-of-origin subtype. Each point represents 1 AOI. Box-and-whisker plots show the median and interquartile range, with whiskers extending to the minimum and maximum values. Statistical analyses are described in the method section.

We performed threshold-independent BayesPrism^52^ deconvolution of CD68^+^ AOI transcriptomes (supplemental Figure 4D). Among macrophage subsets with median abundance >1% across AOIs, TREM2 macrophages were significantly enriched in PCNSL relative to DLBCL (FDR = 0.0199), whereas IL4I1 macrophages showed a nonsignificant trend toward higher abundance in DLBCL (Figure 2E; supplemental Figure 4E).

Given the established role of TREM2 macrophages in lipid handling,^42,46,53^ we next examined lipid-metabolic features in PCNSL macrophages. Within PCNSL CD68^+^ AOIs, an 11-gene lipid-associated macrophage module correlated strongly with a 78-gene TREM2 macrophage module derived from the MoMac-VERSE reference atlas^41^, after excluding genes shared between the two modules (Spearman ρ = 0.86, P < 0.001; Figure 2F; Supplemental Table 4). This correlation exceeded that of all 2000 randomly sampled gene modules of equal size (empirical P < 0.001; supplemental Figure 4F), indicating specificity for the TREM2 macrophage program.

We therefore applied two published macrophage lipid signatures (Figure 2G): one from adipose tissue, where the TREM2-lipid association was originally defined but outside a tumor or CNS context,^46^ and one from glioblastoma, a CNS malignancy with known lipid handling macrophage programs and a tissue context shared with PCNSL.^43^ The former- an adipose tissue macrophage signature- did not differ between PCNSL and DLBCL CD68⁺ regions (P = 0.53; Figure 2H). The second, a GPNMB^high^ lipid-laden macrophage (LLM) signature, was significantly enriched in PCNSL (P = 0.0004; Figure 2J), consistently against both ABC- and GCB-subtype DLBCL (Figure 2K). Within PCNSL, LLM scores were largely restricted to CD68⁺ regions (supplemental Figure 4G).

### LLMs are a conserved, monocyte-derived macrophage (MDM) state distinct from microglia

To determine whether LLM-like features arise from infiltrating MDMs or resident microglia (MG), we analyzed 4 scRNA-seq cohorts (Figure 3A): 2 discovery datasets (Figure 3B) and 2 independent validation cohorts. Because MG has no counterpart in systemic DLBCL, cross-disease comparisons were restricted to MDMs. Major cell types were taken from the annotations provided in each original study, and myeloid subtypes were defined by unsupervised clustering and canonical marker expression, allowing MG to be distinguished from MDMs (supplemental Figure 5A-E). For single-cell analyses, we combined the lipid-associated and GBM-derived gene sets into a single LLM signature (supplemental Methods; Supplemental Table 4).

**Figure 3.**
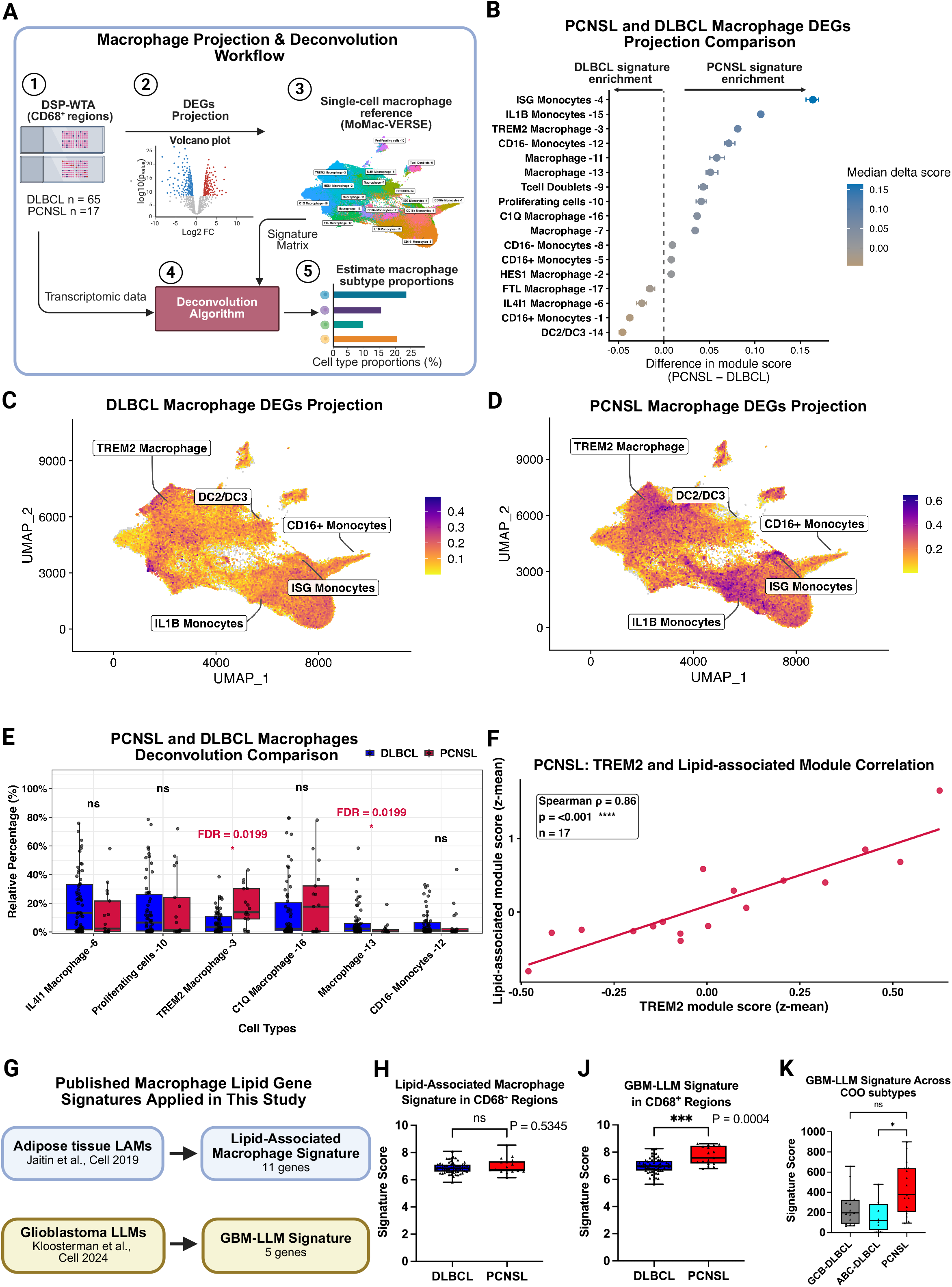
Single-cell RNA sequencing identifies a PCNSL-enriched lipid-laden MDM state. (A) Overview of the scRNA-seq cohorts and analytical workflow. The discovery cohort comprised 8 patients with PCNSL and 17 with DLBCL; validation cohort 1 comprised 13 with PCNSL and 4 with DLBCL; validation cohort 2 comprised 7 with PCNSL and 2 reactive lymph node samples. (B) Uniform manifold approximation and projection (UMAP) of major cell populations in the discovery cohort. (C) LLM signature scores in PCNSL monocyte-derived macrophages (MDMs) and microglia (MG). Upper left, UMAP showing MDM and MG populations; upper right, LLM signature score projected onto the same UMAP; lower panel, comparison of LLM signature scores between MDMs and MG. (D) LLM signature scores across PCNSL MDM clusters. Left, UMAP of MDMs annotated as LLM or non-LLM; right, LLM signature score projected onto the same UMAP. (E) Heatmap of lipid-associated and LLM marker genes in LLMs versus non-LLM MDMs. (F) KEGG over-representation analysis of genes upregulated in LLMs, showing the 5 most enriched pathways (adjusted P < 0.05). (G-H) Patient-level comparison of LLM signature scores between PCNSL MDMs and DLBCL macrophages in the discovery cohort (G) and validation cohort 1 (H). (J) Paired patient-level comparison of LLM signature scores between PCNSL MDMs and MG in validation cohort 1. (K) LLM signature scores across PCNSL MDM clusters in validation cohort 2. Left, UMAP of MDMs annotated as LLM or non-LLM; right, LLM signature scores in LLM and non-LLM MDMs. (L) Heatmap of lipid-associated and LLM marker genes in LLMs versus non-LLM MDMs in validation cohort 2. In panels G, H, and J, each point represents 1 patient. Violin plots show quartiles as dotted lines, with the plot extending to the minimum and maximum values. Statistical analyses are described in the method section.

Single-cell analysis recapitulated the Xenium and DSP-WTA findings: PCNSL had a higher MDM fraction than DLBCL (FDR = 0.019), a comparable T-cell fraction (FDR = 0.672), higher TREM2^+^ and CD163^+^ MDM fractions, and a lower CXCL9:SPP1 ratio consistent across ABC- and GCB-DLBCL (supplemental Figure 5F-J). The functional macrophage genes prioritized in the Xenium and DSP-WTA analyses were expressed predominantly by MDMs rather than microglia (supplemental Figure 6A).

PCNSL MDMs showed higher LLM signature scores than MG at both the cell and patient levels (Figure 3C; supplemental Figure 6B), indicating that the LLM-like program is preferentially associated with infiltrating MDMs. Further unsupervised clustering of PCNSL MDMs identified a TREM2-high subset with the highest LLM signature score (supplemental Figure 6C-E). Over-representation analysis of this subset showed consistent enrichment of lipid-related programs, including cholesterol efflux, fatty acid metabolism, and ether lipid metabolism (supplemental Figure 6F-G); we therefore annotated it as LLMs (Figure 3D).

LLMs expressed lipid-associated and immunoregulatory markers, including TREM2, SPP1, LIPA, and GPNMB, and were enriched for cholesterol-associated, complement and coagulation pathways (Figure 3E-F). Compared with non-LLM macrophages, LLMs exhibited higher lipid-associated, phagocytic, proliferative-associated, and M2-like signatures, but lower M1 and angiogenesis signature scores (supplemental Figure 6H; Supplemental Table 4). LLM abundance varied substantially between patients (supplemental Figure 6J). Compared with DLBCL macrophages, PCNSL MDMs showed higher LLM signature scores at both the patient and cell levels (P = 0.0214 and P < 0.0001, respectively; Figure 3G; supplemental Figure 6K).

These findings were reproduced in two validation cohorts: PCNSL MDMs showed higher LLM signature activity than DLBCL or reactive lymph-node macrophages, and reclustering again identified an SPP1/TREM2/GPNMB/HMOX1/CTSL-expressing LLM-like MDM population (Figure 3H-L; supplemental Figure 7-8).

### LLMs exhibit distinct ligand-receptor crosstalk with T cells

To define how LLMs communicate with other components of the PCNSL TME, we applied LIANA+, an integrative ligand-receptor inference framework, to the PCNSL scRNA-seq discovery cohort.^47^ Global communication analysis identified macrophage/myeloid cells and T cells as the dominant interacting compartments, with macrophage populations showing the strongest overall communication activity and network centrality (Figure 4A; supplemental Figure 9A; Supplemental Table 5).

**Figure 4.**
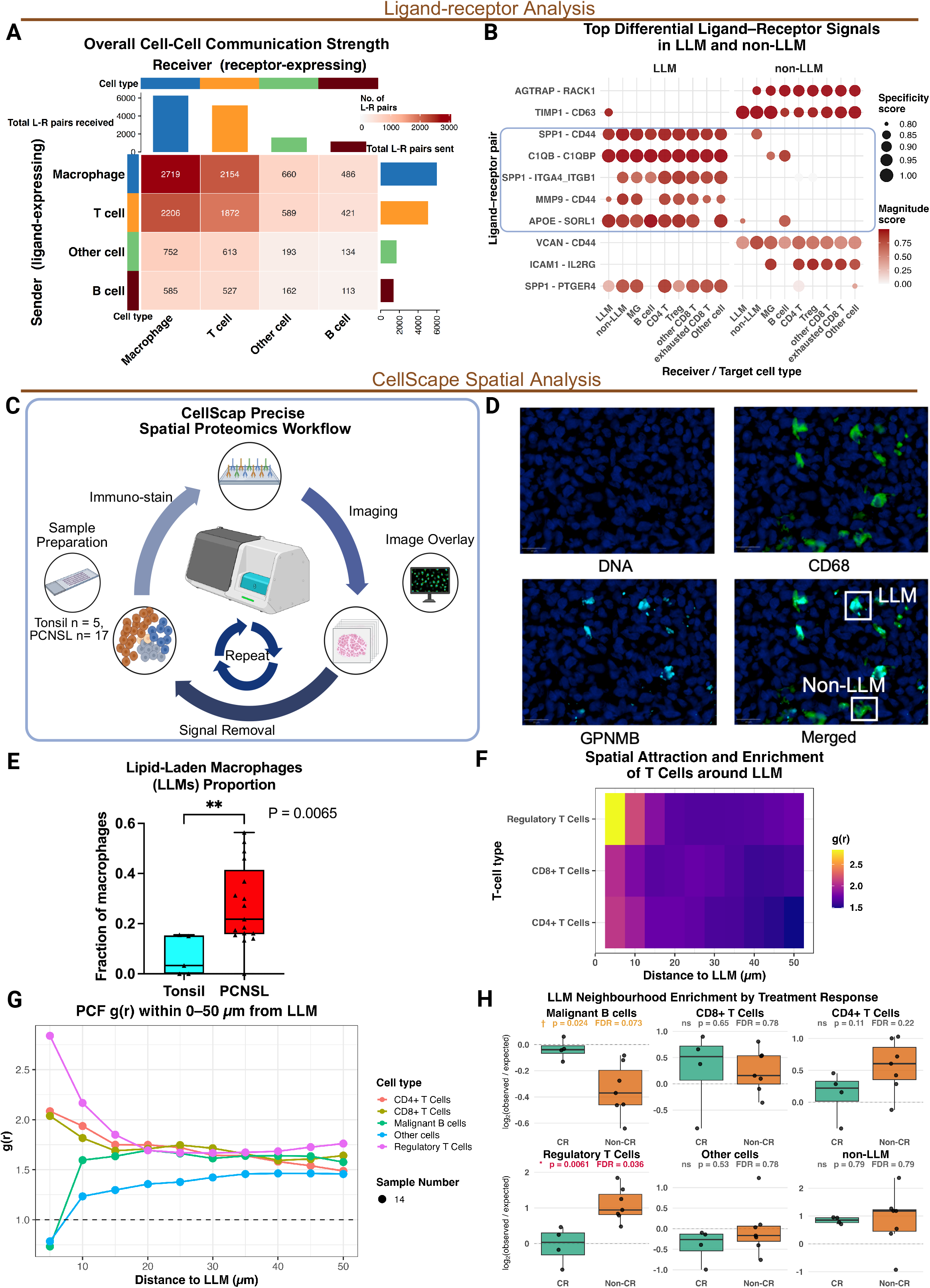
Ligand-receptor inference and spatial proteomic imaging identify LLM-associated communication programs and cellular niches in PCNSL. (A) Heatmap of global ligand-receptor (L-R) interactions among major cell populations in the PCNSL tumor microenvironment. Marginal bar plots show the total number of L-R pairs sent by each cell type (rows, ligand-expressing) and received by each cell type (columns, receptor-expressing). (B) Ligand-receptor pairs with the largest difference in interaction magnitude between LLMs and non-LLM macrophages, shown alongside the corresponding interactions from non-LLM macrophages. Magnitude reflects the estimated strength of an interaction, and specificity reflects how restricted that interaction is to a given pair of cell identities (see in the method section). Dot color represents 1 - magnitude rank and dot size represents 1 - specificity rank; darker and larger dots therefore indicate stronger and more specific inferred interactions. (C) Overview of the CellScape spatial hyperplex proteomic imaging workflow. (D) Representative multiplex image showing the markers used for macrophage phenotyping and identification of LLM-like macrophages. Images were acquired on the CellScape platform and analyzed in QuPath v0.7.0. Scale bar, 20 µm. (E) Proportion of GPNMB⁺CD68⁺ LLM-like macrophages among CD68⁺ macrophages in PCNSL (n = 17) and tonsil controls (n = 5). (F) Cross-type pair-correlation function (PCF) analysis of immune populations around LLM-like macrophages. The heatmap shows PCF-derived enrichment of T-cell populations within LLM-like macrophage neighborhoods across distances of 0 to 50 µm. (G) PCF curves for the same populations, showing g(r) as a function of distance; values above 1 indicate spatial enrichment relative to a random distribution. Curves show median estimates across eligible ROIs from 14 PCNSL samples. (H) Patient-level K-nearest-neighbor analysis of populations neighboring LLM-like macrophages, stratified by treatment response. For each LLM-like macrophage located at least 30 μm from the ROI boundary, the 2 nearest non-LLM cells within 30 μm were identified, and enrichment was expressed as the log_2_ ratio of observed to expected neighbor frequency, with expected frequencies derived from spatially randomized null models. Only ROIs containing at least 20 LLM-like macrophages and at least 20 cells of the partner population were analyzed; 12 of 14 ROIs met these criteria, yielding 11 patients (CR, n = 4; non-CR, n = 7). Measurements from multiple ROIs belonging to the same patient were averaged. Each point represents 1 patient; center lines indicate medians and boxes indicate interquartile ranges. Statistical analyses are described in the method section.

Comparing LLMs with non-LLM macrophages, the interactions showing the largest difference in magnitude included C1QB-C1QBP, SPP1-CD44, SPP1-ITGA4/ITGB1, APOE-SORL1, and MMP9-related axes (Figure 4B), with SPP1, C1QB, and APOE among the most highly expressed ligands in LLMs relative to all other cell populations (supplemental Figure 9B). Ranked by magnitude alone, LLM-derived interactions were directed predominantly toward Tregs, CD4⁺ and CD8⁺ T cells, and myeloid populations (supplemental Figure 9C). Across both rankings, complement, osteopontin, lipoprotein, and matrix-remodeling axes emerged as the dominant LLM-derived signals, implicating them as candidate routes through which LLMs may engage T cells and other myeloid populations.

#### GPNMB^+^ LLMs form regulatory T-cell niches linked to treatment response

To validate LLM-like macrophages at the protein level, we profiled PCNSL tissues (n = 16) and tonsil controls (n = 5) by CellScape using a customized macrophage-focused panel that included the lipid-associated markers GPNMB, TREM2, and CD36 (Figure 4C; supplemental Figure 9D). Among these, GPNMB gave the most consistent staining and is the defining marker of the GBM-derived LLM signature; we therefore defined GPNMB^+^CD68^+^ cells as LLM-like macrophages for spatial analysis (Figure 4D). These cells were significantly more abundant in PCNSL than in tonsil controls (P = 0.0065; Figure 4E). Pair-correlation analysis showed that LLM-like macrophages were spatially enriched near T cells, particularly within 10 μm, with FOXP3^+^ Tregs showing the strongest local enrichment (Figure 4F-G), concordant with the ligand-receptor inference.

We next examined whether LLM-associated neighborhoods differed by treatment response. All patients received frontline high-dose methotrexate-based chemotherapy, with or without consolidative autologous stem cell transplantation or radiotherapy, and were classified at the end of first-line treatment as complete response (CR, n = 4) or non-CR (n = 7). Non-CR patients showed a numerically higher LLM fraction among macrophages, although this comparison did not reach significance (supplemental Figure 9E). In non-CR cases, FOXP3⁺ Tregs were found within 30 μm of LLM-like macrophages about twice as often as expected by chance (P = 0.0061, FDR = 0.036), a finding which was not noted in CR cases. A reciprocal trend was observed for malignant B cells (P = 0.024, FDR = 0.073). These differences were stable across neighbor counts and distance thresholds, and the Treg association was attenuated but retained when LLM identity was permuted within the macrophage compartment (Δ 1.14 to 0.73; P = 0.012), indicating that it is not fully explained by shared niche occupancy; the malignant B-cell trend did not persist under this null (supplemental Figure 9F-G). Neither CD4^+^ nor CD8^+^ T cells differed between response groups.

Together, the ligand-receptor and spatial analyses converge on the same relationship that LLMs signal preferentially to T cells, and Tregs accumulate in their immediate vicinity.

### An immunocompetent MCD-driven model shows LLMs are locally induced by CNS lymphoma

The data from PCNSL patients establish LLMs as a recurrent immunosuppressive feature, but do not determine whether these cells are induced by lymphoma within the CNS or reflect a pre-existing brain macrophage population. We developed the first immunocompetent, syngeneic, MCD-genotype-driven model, in which murine MCD-like DLBCL cells (MCD_P2), derived from previously published mouse models,^54–57^ were implanted intracranially into wild-type C57BL/6 mice, allowing matched comparison of the tumor-bearing side (TS) and contralateral non-tumor side (NTS) (Figure 5A). Tumors remained CNS-restricted without systemic dissemination.

**Figure 5.**
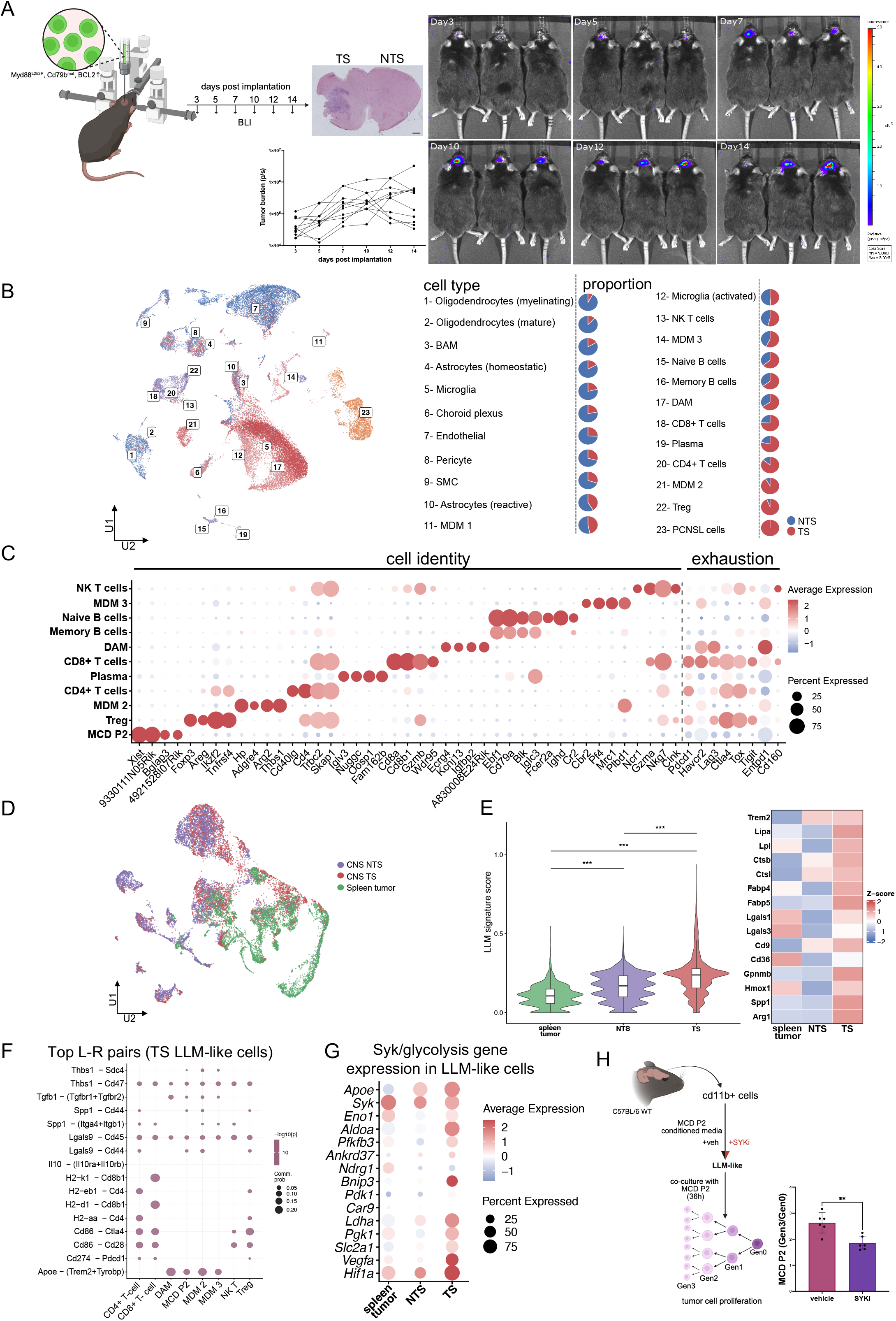
An immunocompetent, MCD-genotype-driven model of PCNSL shows that lipid-laden macrophages are locally induced by CNS lymphoma and are reversible by SYK inhibition. (A) Schematic of the immunocompetent syngeneic PCNSL model. Murine MCD-DLBCL cells (MCD_P2; Myd88L252P, Cd79b-mutant, Bcl2-overexpressing) were implanted into the left cerebral hemisphere of wild-type C57BL/6 mice, enabling matched within-animal comparison of the tumor-bearing side (CNS TS) and contralateral non-tumor side (CNS NTS); parallel splenic implants of the same cells provided a systemic (extracranial) comparator driven by identical oncogenes. Tumors remained CNS-restricted without systemic dissemination. (B) UMAP of the 22 CNS tumor-microenvironment cell types resolved by single-cell RNA-seq. See also Fig. S10A. Pie charts indicate the proportional composition of each cell type in CNS TS and CNS NTS. (C) Expression of exhaustion (*Pdcd1*, *Havcr2*, *Lag3*, *Ctla4*, *Tox*, *Tigit*) and cytotoxicity (*Gzma*, *Nkg7*) markers across CNS TS CD8^+^, CD4^+^, and regulatory T-cell populations, indicating an immunosuppressive, inflammatory lymphoma microenvironment. (D) UMAP of the myeloid cells identified in CNS TS, CNS NTS and spleen tumor. (E) LLM signature score and individual LLM gene expression across regional myeloid populations, showing the anatomic gradient CNS TS > CNS NTS > splenic tumor (violin plot, signature score; heatmap, individual genes). This gradient establishes that the LLM state is acquired locally within the CNS in response to lymphoma rather than reflecting a systemic myeloid response to MCD-driven disease. See also in Figure S10B-E. (F) Ligand-receptor analysis identifying Apoe-(Trem2+Tyrobp) signaling among CNS TS myeloid (DAM, MDM2, MDM3) and tumor (MCD_P2) populations, upstream of spleen tyrosine kinase (SYK). (G) Expression of *Syk* and downstream glycolytic and HIF1α-associated genes (*Pfkfb3*, *Ldha*, *Aldoa*, *Eno1*, *Pgk1*, *Pdk1*, *Hif1a*, *Bnip3*, Ndrg1, *Ankrd37*, *Car9*) across CNS TS LLM-like clusters (MDM2), consistent with SYK-driven glycolytic reprogramming. (H) Functional assessment of lymphoma-conditioned LLM-like myeloid cells. Primary CD11b^+^ brain myeloid cells were cultured in MCD_P2-conditioned medium with vehicle or the SYK inhibitor BAY 61-3606, then co-cultured with CellTrace Violet-labeled MCD_P2 lymphoma cells for 36h. Representative CellTrace Violet dilution profiles and Gen3/Gen0 ratio quantification (1.76 vs 2.67; P = 0.0087) show that SYK inhibition within LLM-like cells alone reduces lymphoma-cell proliferation, demonstrating that the tumor-supportive activity of these cells is pharmacologically reversible.

Single-cell profiling resolved 23 cell types in the CNS (Figure 5B; supplemental Figure 10A). Relative to NTS, the TS showed expansion of activated and disease-associated microglia (DAM), three MDM populations (MDM1, MDM2, MDM3), and exhausted, cytotoxic CD8^+^/CD4^+^ T cells and Tregs expressing *Pdcd1*, *Lag3*, *Ctla4*, *Tox*, and *Tigit* (Figure 5C), recapitulating the immunosuppressive inflammatory microenvironment of human PCNSL. Of these, the myeloid population MDM2 expressed high LLM genes, mirroring human LLMs (supplemental figure 10A).

Critically, in a three-way comparison of all myeloid cells from the CNS TS, CNS NTS and splenic tumor driven by the same oncogenes, the LLM signature was highest in CNS TS (Figure 5D-E). Unsupervised clustering of myeloid cells from all CNS and spleen tumor samples identified 24 distinct clusters (supplemental figure 10B). Of these, cluster C7 (supplemental Figure 10C-D) was identified as transcriptionally most similar to the MDM2 LLM-like myeloid population. C7 from the CNS TS showed the highest LLM signature score compared to CNS NTS and spleen tumor (supplemental Figure 10E). This anatomic gradient establishes that the LLM state is acquired locally within the CNS in response to lymphoma, rather than reflecting a systemic myeloid response to MCD-driven disease.

### SYK inhibition reverses the tumor-supportive activity of LLM-like macrophages

Ligand-receptor analysis in the murine model recovered Spp1-mediated axes directed toward T cells, mirroring the human inference, and additionally identified Apoe-Trem2/Tyrobp signaling between LLM-like macrophages and tumor cells (Figure 5F).

Because ApoE-TREM2/TYROBP acts upstream of spleen tyrosine kinase (SYK), we examined SYK signaling in CNS TS LLM-like cells and found elevated expression of *Syk* together with downstream glycolytic and HIF1α-associated genes (*Pfkfb3*, *Ldha*, *Aldoa*, *Eno1*, *Pgk1*, *Pdk1*, *Hif1a*, *Bnip3*, *Ndrg1*, *Ankrd37*, *Car9*; Figure 5G), consistent with SYK-driven glycolytic reprogramming of myeloid cells.^58,59^

We therefore tested whether SYK inhibition can alter LLM-like state in myeloid cells and its tumor-supportive activity. CD11b^+^ brain myeloid cells cultured in MCD_P2-conditioned medium acquired a TREM2^+^CD163^+^ phenotype, which was reduced by the SYK inhibitor BAY 61-3606 (SYKi; supplemental Figure 10F). In co-culture, LLM-like cells pre-treated with SYKi reduced proliferation of CellTrace-labeled MCD_P2 PCNSL cells (Gen3/Gen0 ratio, 1.76 vs 2.67; P = 0.0087; Figure 5H). Inhibiting SYK within LLM-like cells alone was therefore sufficient to blunt lymphoma-cell proliferation, providing functional evidence that LLM-mediated tumor support is pharmacologically reversible.

## Discussion

PCNSL is usually classified as DLBCL, with genetic and transcriptomic features similar to MCD-subtype DLBCL.^60^ However, it exhibits CNS restriction and distinct clinical behavior, which are incompletely explained by tumor-intrinsic features alone. Combining spatial multi-omics across PCNSL and systemic DLBCL with the first immunocompetent, MCD-like-genotype-driven mouse model of PCNSL, we identify lipid-laden macrophages (LLMs) as a CNS-enriched, lymphoma-induced myeloid state that occupies immunosuppressive niches and supports tumor growth. The concordance between human tissue and a genetically faithful animal model, including the anatomic specificity of LLM induction and the reduction of their tumor-supportive activity by SYK inhibition, suggests the LLM program to be mechanistically important and a therapeutic target in CNS lymphoma.

Unlike several other genetic and transcriptomic features that are shared between PCNSL and systemic DLBCL, this LLM program is a clear distinction between the two entities. Of interest, it overlaps with lipid-laden TAM states described in glioblastoma and other CNS inflammatory settings, where macrophages acquire lipid-rich, myelin-scavenging phenotypes.^43,61^ Although previous single-cell studies characterized the PCNSL TME, including comparisons with systemic DLBCL, they generally grouped MDMs and microglia within broad myeloid compartments.^15,21,62^ Our analysis instead resolves LLMs by cellular origin, spatial context, and disease site. LLM signatures were enriched in PCNSL MDMs and FOXP3⁺ Treg niches and, in the first immunocompetent model, are preferentially induced at intracerebral tumor sites relative to contralateral brain and genotype-matched splenic tumors. This pattern is distinct from the broadly similar adaptive immune profiles in intracerebral and intrasplenic lymphomas noted in a prior murine cell implantation model which lacked PCNSL specific gene alterations.^63^ In keeping with the human data, restriction of the LLM program to lymphoma-involved brain regions in the MCD model further supports local, lymphoma-associated reprogramming rather than nonspecific CNS residence.

We propose that LLMs arise through the convergence of lymphoma-derived signals with the lipid-rich, myelin-containing CNS milieu. Local lymphoma growth may expose recruited MDMs to cholesterol-rich myelin debris and tumor-derived cargo. Recognition of these ligands through the APOE-TREM2/TYROBP-SYK axis identified in our syngeneic model could drive their uptake, establishing a feed-forward loop in which lipid acquisition further reinforces TREM2 signaling. SYK activation may also promote glycolytic reprogramming, consistent with the coordinated upregulation of Syk and glycolytic/HIF-1α-associated genes in MDMs from CNS tumor sites. Enhanced glycolysis could, in turn, reinforce macrophage immunoregulation through lactate-driven histone lactylation and IL-10 induction, as demonstrated in glioblastoma MDMs.^64^ Prior in vivo evidence that PCNSL-associated macrophage polarization is pharmacologically reversible further supports the plasticity of this myeloid state.^65^ Consistent with this concept, SYK inhibition reduced LLM-mediated tumor support ex vivo, providing functional evidence for a coupled lipid-glycolytic program. Together, these findings nominate TREM2-SYK dependence and APOE-SORL1-mediated lipid transfer as testable mechanisms of lymphoma support. The latter has precedent in glioblastoma, where myelin uptake enables macrophage-to-tumor lipid transfer.^43^

This lipid-remodeled state may establish a Treg-permissive, tumor-supportive niche. Macrophage-derived SPP1 can engage CD44 and integrin receptors on T cells,^66,67^ while SPP1⁺ macrophages promote CD137⁺ Treg maturation through SPP1-CD44-NF-κB1 signaling in colorectal cancer tumor-draining lymph nodes.^68^ This precedent may explain the inferred SPP1-CD44 and SPP1-ITGA4/ITGB1 interactions in LLM-rich regions, which could facilitate Treg maturation or retention.

Consistent with this, prior PCNSL studies on the microenvironment have described immunosuppressive macrophages and T-cell exhaustion and microglial phenotypes, and macrophage plasticity, supporting the rationale for macrophage-targeted strategies to facilitate macrophage reprogramming and T-cell reinvigoration.^17,25–29,69,70^ Spatially resolved work has extended these observations to tissue architecture, describing perivascular T-cell and chemokine organisation,^71^ global microenvironmental archetypes,^16^ with macrophage-centred proximities and pericyte-derived CCL19 carrying prognostic information beyond immune-cell density alone.^37,49^ What has remained unresolved, however, is the precise identity and functional state of the macrophages populating these architectures, and how this compartment differs from that of nodal DLBCL, a gap that has kept macrophage-targeted therapeutic rationale broad rather than actionable. By resolving the macrophage compartment by ontogeny and metabolic state, we identify a lipid-remodeled, monocyte-derived population occupying a discrete, Treg-enriched microniche associated with chemotherapy response. This finding moves beyond a general case for macrophage-directed therapy toward a specific, spatially defined cellular target: rather than a compositional category, we define a molecularly distinct macrophage state and its niche as a candidate point of therapeutic intervention. More broadly, our data suggest that the immune architectures described in prior spatial studies may themselves comprise functionally heterogeneous local niches not resolved by immune-cell density or broad lineage markers alone and that resolving these niches is necessary to translate microenvironmental biology into precise therapeutic strategies.

Beyond SYK, our findings of the LLM population in PCNSL nominate other potentially actionable nodes. Recent preclinical studies have described strategies targeting TREM2⁺ TAMs, including IL-2 myeloid-targeted immunocytokines and natural killer (NK)/T cell enhancers (MiTEs), IL-12-armored TREM2 CAR T cells, and direct inhibition, which reprogrammed suppressive myeloid cells, enhanced cytotoxic immunity, and improved responses to PD-1 blockade.^72–74^ Anti-GPNMB CAR T cells have likewise shown activity against GPNMB⁺ tumor and immunosuppressive myeloid cells.^75^

Several limitations should be noted. Our MDM-versus-microglia discrimination relies primarily on single-cell RNAseq derived annotation. Protein-level validation focused on GPNMB, with the sample sizes throughout this study limited by the rarity of PCNSL. Multiplexed protein-level comparisons between PCNSL and systemic DLBCL are warranted to supplement the transcriptomic findings. While the results are in keeping with human spatial and transcriptomic data, the mechanistic studies were performed in a syngeneic mouse model as patient-derived xenografts lack macrophage-T cell interplay.

In summary, PCNSL contains a lipid-laden, immunoregulatory MDM program that is transcriptionally distinct from microglia and enriched within the CNS lymphoma microenvironment. Using the first immunocompetent MCD-like-driven PCNSL mouse model, we further demonstrate that this program is induced locally by lymphoma and is amenable to pharmacological modulation. Together, these findings position macrophage immunosuppression and lipid remodeling as central features of the PCNSL microenvironment and identify LLMs as a candidate therapeutic target.

## Acknowledgement

A.D.J. is supported by the Singapore Ministry of Health’s National Medical Research Council (NMRC) Clinician Scientist Award (MOH-00715-00; CSASI24jul-006). Research in ADJ’s laboratory is funded by the Core Grant from the Cancer Science Institute of Singapore, National University of Singapore through the National Research Foundation, Singapore, and the Ministry of Education under the Research Centres of Excellence initiative.

M.L. was supported by the National Natural Science Foundation of China (Grant No. 82500239), Chongqing Municipal Natural Science Foundation (Grant No. CSTB2024NSCQ-MSX1080), The 76th Batch of the China Postdoctoral Science Foundation General Program (2024M763884), Chongqing Municipal Medical Young Top Talent Program (YXQN2025027), and Chongqing Municipal Postdoctoral Special Funding Program (2312013533263948).

A.R. was supported by NIH 5P30CA046592, University of Michigan Rogel Cancer Center Support Grant.

L.C. was supported by the National Cancer Institute P01 CA272295. BCU SCOR 7027-23 and BCU TRP 6695.

HCR was supported by the German Research Foundation (DFG) (SFB1399- grant no. 413326622, SFB1430 - grant no. 424228829, and SFB1530 - grant no. 455784452, RE 2246/17-1 - 553375105, SFB1752 - grant no. pending), the German Cancer Aid (1117240, 70113041, the Mildred Scheel Nachwuchszentrum Grant 70113307, the TACTIC consortium as part of the Preclinical Drug Development Program preCDD and an Excellence Funding Program grant, the TransOnk consortium ERASE), the German Ministry of Education and Research (BMBF e:Med Consortium InCa, grants 01ZX1901, 01ZX2201A and the DKTK grant BOOST-CAR-T) and the CANcer TARgeting (CANTAR) project NW21-062 “Netzwerke 2021” an initiative of the Ministry of Culture and Science of the State of North Rhine-Westphalia.

During the preparation of this work, the authors used Claude (Opus 4.8, Anthropic) and ChatGPT (OpenAI, San Francisco, CA, USA; accessed 23 July 2026) to improve the clarity and readability of the writing in the manuscript-particularly for the Introduction and Discussion sections. All AI-generated suggestions were reviewed and edited by the corresponding author (A.D.J) and the bioinformatician (L.H.). No AI tools were used in study design, sequencing analysis, variant interpretation, or clinical correlations. The authors take full responsibility for the content of this article.

## Authorship Contributions

A.D.J., C.Tr., and L.C. conceived the study and supervised the project. L.H. designed the analytical framework, performed the spatial transcriptomic, single-cell transcriptomic, spatial proteomic, and computational analyses, interpreted the data, and wrote the initial manuscript draft. M.L. performed the digital spatial profiling of PNCSL and DLBCL, and S.S contributed to all the bioinformatic and spatial analysis of the human data. S.J. and M.V.R. established the murine lymphoma models, generated experimental data, and contributed to murine data collection and preprocessing. C.L.T., S.S.-S.H., S.-B.N., S.D.M., L.P., Y.B., and J. contributed clinical samples, pathology review and interpretation, and clinical data curation. R.S. and A.T. performed Xenium data processing and cell annotation, with input from A.R. O.B, D.J.S, J.B and F.H performed and analyzed the Cellscape data. N.Z. and F.R.M. contributed to mouse data acquisition. G.K. and H.C.R. contributed to the development of the murine cell line. Z.Y.C.O., S.C.N.T., W.X.C.L., C.Ti., P.J., Y.P., J.B., O.B., E.T.K., and other co-authors contributed to data interpretation, study design discussions, and critical revision of the manuscript. M.X. and J.C. contributed to the review and robustness assessment of the single-cell analyses. F.H., B.J.L., D.J.-S., Q.P.-H., Z.H., M.B., N.T., C.K., and M.P. provided key resources, technical support, and scientific input. All authors reviewed, edited, and approved the final manuscript.

## Conflicts of Interest Disclosure

Anand D. Jeyasekharan has received consultancy fees from DKSH/BeiGene, Roche, Gilead, Turbine Ltd, AstraZeneca, Antengene, Janssen, MSD, IQVIA, and KYAN Technologies, and research funding from Janssen and AstraZeneca; all of which are unrelated to this work.

A.R. serves as a member of Voxel Analytics, LLC, and consults for Telperian. He also serves as a faculty advisor for TCS Ltd. and as the Satish Dhawan Visiting Chair Professor at Indian Institute of Science Bangalore.

HCR received consulting and lecture fees from Abbvie, Roche, KinSea, Vitis, Cerus, Lilly, Novartis, Takeda, AstraZeneca, Vertex, and Merck. HCR received research funding from AstraZeneca and Gilead Pharmaceuticals. HCR is a co-founder of CDL Therapeutics GmbH.

OB, DJS, JB and FH are current employees of Bruker Corporation and may hold stock and/or stock options in the aforementioned company. Bruker Corporation is the developer of the CellScape technology used in this study.

## Tables

Attached file ‘Supplementary Table 1.xlsx’.

**Supplemental Figure 1. Xenium profiling resolves canonical marker expression and cell-type composition across TMA cores.**

(A)Dot plot of canonical marker expression across annotated cell types. Dot size indicates the proportion of cells expressing each gene, and dot color indicates mean expression. (B) Cell-type composition of all individual TMA cores at the lineage level. Each column represents 1 TMA core.

**Supplemental Figure 2. Xenium profiling of immune-cell composition in PCNSL and DLBCL.**

(A) Representative TMA core showing selected canonical markers from the 380-gene Xenium Immuno-Oncology panel. Images were acquired on the Xenium Analyzer (PN-1000481; 10x Genomics), which performs automated sequential fluorescence imaging, transcript decoding, and onboard analysis, and were visualized in Xenium Explorer version 4.0. (B-C) Cell-type composition of individual DLBCL samples at the broad lineage level (B) and at finer subtype resolution (C). Each column represents 1 TMA core. (D) Pan-T-cell proportions in PCNSL (n = 17 cores) and DLBCL (n = 171 cores). (E) CD4:CD8 T-cell proportion ratios in PCNSL and DLBCL. In panels D and E, each point represents 1 TMA core; box-and-whisker plots show the median and interquartile range, with whiskers extending to the minimum and maximum values. Statistical analyses are described in the method section.

**Supplemental Figure 3. Quality control of GeoMx DSP WTA data.**

(A) Gene-level quality control. The dashed line indicates the log_2_ CPM detection threshold (4.08); genes falling below this threshold in at least 90% of AOIs would be removed. No genes met this criterion in the present dataset. (B) AOI-level quality control based on nuclei count; AOIs with 20 or fewer nuclei were excluded. All 17 PCNSL and 62 of 64 DLBCL CD68^+^ AOIs were retained. (C-D) Relative log expression across all AOIs before (C) and after (D) TMM normalization. (E-F) Heatmaps of the top PCNSL-upregulated genes before (E) and after (F) TMM normalization, showing that expression trends were preserved. (G) Representative morphology images showing CD68-AF594, CD20-AF647, CD3-AF532, and SYTO 13 nuclear staining. Images were acquired on the GeoMx Digital Spatial Profiler (NanoString) using GeoMx DSP software version 2.4.0.421. ROIs were selected under pathologist guidance, and fluorescence-channel thresholds were adjusted to generate CD68^+^, CD3^+^, and CD20^+^ masks. (H) Cumulative distributions of lineage-marker gene expression across AOI masks. (J) T-cell density in PCNSL versus DLBCL CD3^+^ AOIs. (K) Running enrichment plots for the 3 most significantly enriched Hallmark gene sets among PCNSL-upregulated genes.

**Supplemental Figure 4. MoMac-VERSE signature projection and BayesPrism deconvolution of CD68^+^ AOIs.**

(A) UMAP of the MoMac-VERSE macrophage reference atlas. (B) Dot plot showing expression of PCNSL- and DLBCL-enriched differentially expressed genes across macrophage states in the MoMac-VERSE reference. Dot size indicates the proportion of cells expressing each gene, and dot color indicates mean expression. (C) Delta module scores across macrophage states, calculated as the PCNSL signature score minus the DLBCL signature score. (D-E) BayesPrism-estimated macrophage-state proportions in PCNSL and DLBCL CD68^+^ AOIs, shown as the composition of individual samples (D) and as group-wise comparisons of individual macrophage states (E). Both panels include states below the 1% median abundance threshold applied in Figure 2E. (F) Specificity of the correlation between TREM2-macrophage and lipid-associated module scores (observed ρ = 0.86; empirical P < 0.001). The histogram shows Spearman correlation coefficients between the lipid-associated module score and 2000 randomly sampled gene modules of equal size (78 genes) drawn from the WTA panel; the vertical line indicates the observed correlation with the TREM2 macrophage module. (G) GBM-derived GPNMB^high^ LLM signature scores across CD3^+^, CD20^+^, and CD68^+^ AOIs in PCNSL. Each point represents 1 AOI. Box-and-whisker plots show the median and interquartile range, with whiskers extending to the minimum and maximum values. Statistical analyses are described in the method section.

**Supplemental Figure 5. Myeloid annotation and immune composition in the discovery scRNA-seq cohort.**

(A) UMAP of annotated myeloid populations in PCNSL. (B) Dot plot of canonical marker expression across myeloid populations. (C) UMAP of annotated macrophage (TAM) populations, re-clustered from the TAM compartment shown in panel A. (D) Dot plot of canonical marker expression across macrophage populations. In panels B and D, dot size indicates the proportion of cells expressing each gene, and dot color indicates mean expression. (E) Correspondence between author-provided annotations from the original publications and the harmonized annotations used in this study. (F-G) Patient-level proportions of MDMs or macrophages (F) and T cells (G) in PCNSL and DLBCL. (H) Patient-level comparison of CD163^+^ and TREM2^+^ macrophage fractions and CXCL9:SPP1 ratios between PCNSL MDMs and DLBCL macrophages. (J) Corresponding comparisons between PCNSL MDMs and ABC- or GCB-subtype DLBCL macrophages. In panels F-J, each point represents 1 patient; box-and-whisker plots show the median and interquartile range. Statistical analyses are described in the method section.

**Supplemental Figure 6. The LLM signature defines a lipid-enriched MDM subset in the PCNSL discovery cohort.**

(A) Patient-level heatmap comparison of macrophage functional gene expression in PCNSL MDMs and microglia. Values are row-scaled z scores. (B) Paired patient-level comparison of LLM signature scores between PCNSL MDMs and microglia. (C) LLM module scores across unsupervised PCNSL MDM clusters (resolution = 0.3). (D) UMAP of LLM signature scores (upper) and unsupervised MDM clustering (lower). (E) Heatmap of the top 10 marker genes for each PCNSL MDM cluster. (F-G) Top enriched Gene Ontology biological process (F) and KEGG (G) pathways in MDM cluster 4, which showed the highest LLM signature score and high TREM2 expression. (H) Scores for curated macrophage functional signatures in LLM and non-LLM macrophages, comprising lipid-associated and LLM-defining gene sets, phagocytosis, proliferative-region-associated macrophage (PAM), M1-like, M2-like, and angiogenesis programs. Signature gene sets are listed in Supplemental Table1. (J) LLM and non-LLM fractions across individual patients with PCNSL. (K) Cell-level comparison of LLM signature scores between PCNSL MDMs and DLBCL macrophages. In panel B, each point represents 1 patient. Box-and-whisker and violin plots show the median and interquartile range. Statistical analyses are described in the method section.

**Supplemental Figure 7. LLM signature enrichment in PCNSL macrophages in validation cohort 1.**

(A) UMAP of all cells from PCNSL and DLBCL samples colored by disease. (B) UMAP shown separately for PCNSL (left) and DLBCL (right), colored by annotated major cell types. (C) Heatmap of top marker genes for each annotated cell type in PCNSL. (D-E) Cell-level comparison of LLM signature scores between PCNSL MDMs and DLBCL macrophages (D) and between PCNSL MDMs and microglia (E). Violin plots show quartiles as dotted lines, with the plot extending to the minimum and maximum values. Statistical analyses are described in the method section.

**Supplemental Figure 8. LLM-like macrophage features in PCNSL in validation cohort 2.**

(A) UMAP shown separately for PCNSL (left) and reactive lymph node (right), colored by annotated major cell type. (B) Dot plot of canonical marker expression across PCNSL myeloid populations. Dot size indicates the proportion of cells expressing each gene, and dot color indicates mean expression. (C) UMAP of annotated PCNSL myeloid populations (left) and LLM signature score projected onto the same UMAP (right). (D) UMAP of unsupervised PCNSL MDM clusters (resolution, 0.6). (E) LLM module scores across unsupervised MDM clusters. (F) Dot plot of the 5 most enriched KEGG pathways in LLMs. Dot size indicates the number of genes in each pathway, and dot color indicates −log₁₀(adjusted P). (G) Fine cell-type composition of individual patients with PCNSL. (H-J) LLM signature scores in PCNSL MDMs compared with reactive lymph node macrophages (H) and with microglia (J), shown at the cell level (left) and patient level (right). In patient-level comparisons, each point represents 1 patient. Statistical analyses are described in the method section.

**Supplemental Figure 9. Ligand-receptor inference, spatial imaging workflow, and robustness of neighborhood analyses in PCNSL.**

(A) Network centrality of major PCNSL cell populations based on inferred ligand-receptor communication in the discovery cohort. The x-axis shows outgoing centrality, defined as the number of inferred interactions in which a cell type acts as the sender (ligand-expressing), and the y-axis shows incoming centrality, defined as the number of interactions in which it acts as the receiver (receptor-expressing). (B) Dot plot showing expression of key ligand genes across PCNSL cell types. Dot size indicates the proportion of cells expressing each gene, and dot color indicates mean expression. (C) Heatmap of the top 10 ligand-receptor pairs sent by LLMs to each recipient cell type, ranked by interaction magnitude (defined in Figure 4B), with color indicating magnitude score. (D) Overview of the TMA image-processing workflow, including image acquisition, cycle alignment, single-cell segmentation, marker thresholding, phenotype assignment, and population quantification. (E) Proportion of GPNMB^+^CD68^+^ LLM-like macrophages among CD68⁺ macrophages in patients with PCNSL, stratified by treatment response. The same ROI inclusion criteria as in Figure 4H were applied. Box plots show the median and interquartile range. (F-G) Sensitivity analyses for the neighborhood enrichment shown in Figure 4H. (F) Robustness to spatial parameter choice, shown for regulatory T cells (left) and malignant B cells (right). Each cell of the heatmap gives the difference in mean log_2_(observed/expected) enrichment between non-CR and CR patients for one combination of k, the number of nearest non-LLM neighbors considered, and r_cap, the maximum distance at which a neighbor is counted. Warmer colors indicate stronger enrichment in non-CR cases and cooler colors the reverse. Asterisks denote nominal P < 0.05, uncorrected for multiple comparisons. The k axis is categorical. Values shown use a minimum of 20 LLM per ROI; thresholds of 10 and 30 gave concordant results. (G) Comparison of null models used to define expected neighborhood composition. Complete spatial randomness (CSR) redistributes all cells uniformly within the ROI and is the null applied in Figure 4H. The macrophage swap null permutes LLM and non-LLM labels within the macrophage compartment while leaving the positions and identities of all other cells unchanged, thereby distinguishing LLM-specific proximity from shared niche occupancy; LLM comprised 15-58% of macrophages in contributing ROIs (median 34%). The toroidal shift null displaces LLM cells by a random cyclic translation, preserving LLM clustering while breaking spatial registration with partner cell types. Bars show the difference in mean log_2_(observed/expected) enrichment between response groups, colored by significance. A global label permutation gives results numerically indistinguishable from CSR and is not shown. The non-LLM compartment is omitted as it constitutes the label permuted under the swap null. Statistical analyses are described in the Methods section.

**Supplemental Figure 10. Murine lymphoma models recapitulate a CNS-specific human LLM-like MDM state which can be functionally modulated by SYK inhibition.**

(A) Expression of canonical marker genes used to identify cell populations within the PCNSL microenvironment. (B) UMAP of myeloid cells isolated from MCD_P2-implanted spleen tumors and from CNS tumor-associated (CNS TS) and CNS non-tumor-associated (CNS NTS) tissue, by unsupervised clustering into 24 transcriptionally distinct clusters (C0-C23) and (C) by per-cell LLM (lipid-laden macrophage) signature score. (D) Stacked bar plot showing the proportional distribution of each myeloid cluster across spleen tumor, CNS TS, and CNS NTS tissue. Clusters C12, C14, C18, C22, and C23 contained too few cells to reliably determine proportions and were omitted from the graph. Cluster C7, identified as the transcriptional nearest neighbor to the MDM2 LLM-like myeloid population in CNS TS, is indicated on the graph. (E) Heatmap of z-scored LLM gene signature expression across individual myeloid clusters (C0-C21) showing highest overall expression in CNS TS cluster C7. (F) Schematic representing TREM2/TYROBP-SYK signaling driving the polarization of CD11b^+^ cells into LLM-like cells and inhibition of that polarization by BAY 61-3606 (SYKi). Quantification of the percentage of TREM2^+^CD163^+^ cells over total CD11b^+^ cells under control, conditioned-media+vehicle (cm+veh), and conditioned-media+SYKi (cm+SYKi) conditions (right; *p < 0.05, ns = not significant), with representative flow cytometry plots shown below for each condition.

## Figures

See in separate attached files.

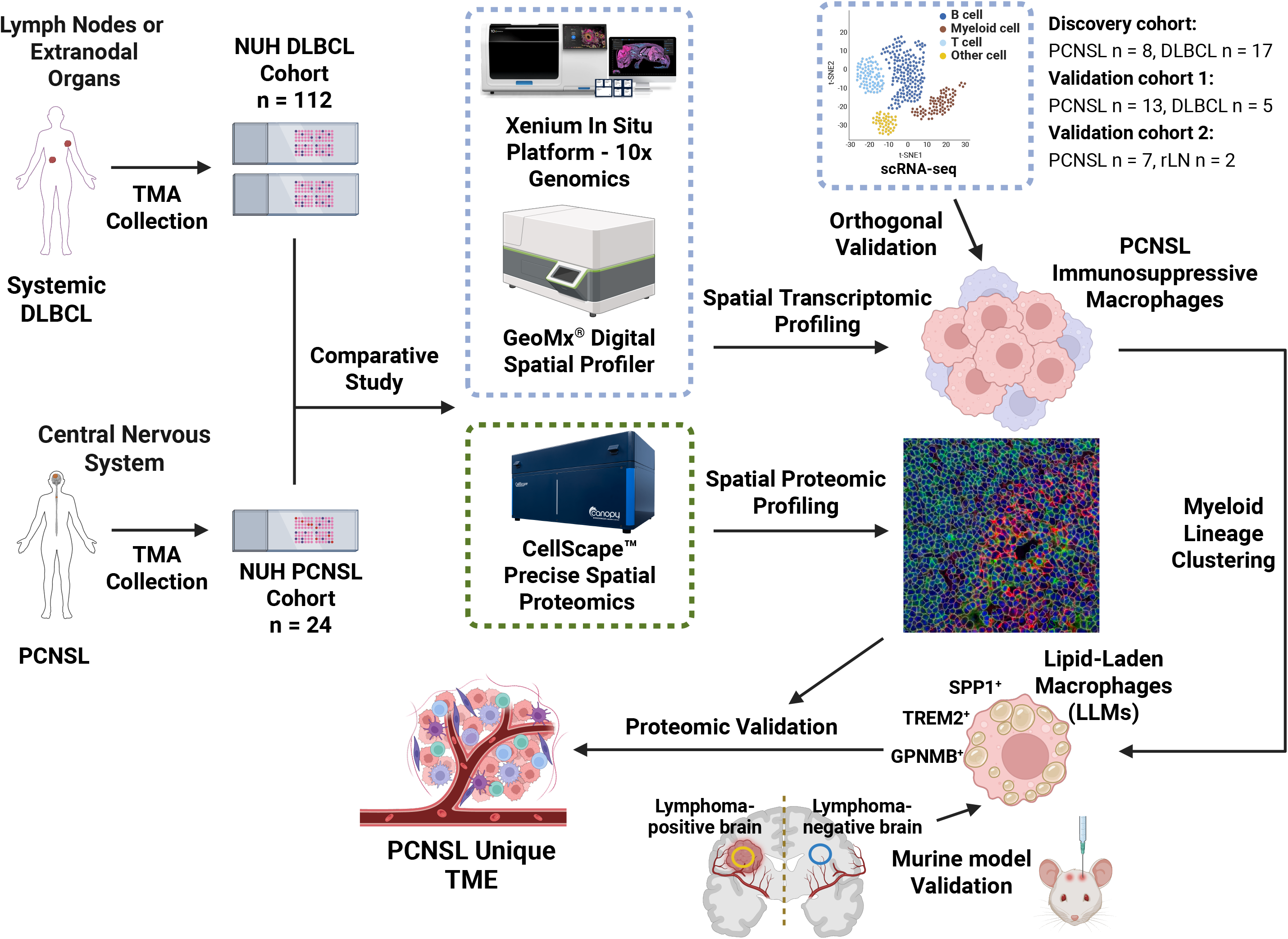

## Notes

### Summary of Updates

Introduce the first immunocompetent PCNSL syngeneic model and functional analysis. Adding new collaborations and funding parties; Figures revised and rearranged; Supplemental files updated.

