## Supplementary figures and images for "A lipid-laden macrophage niche drives immunosuppression in primary central nervous system lymphoma"

### Supplementary FigureS1

#### Slide 1
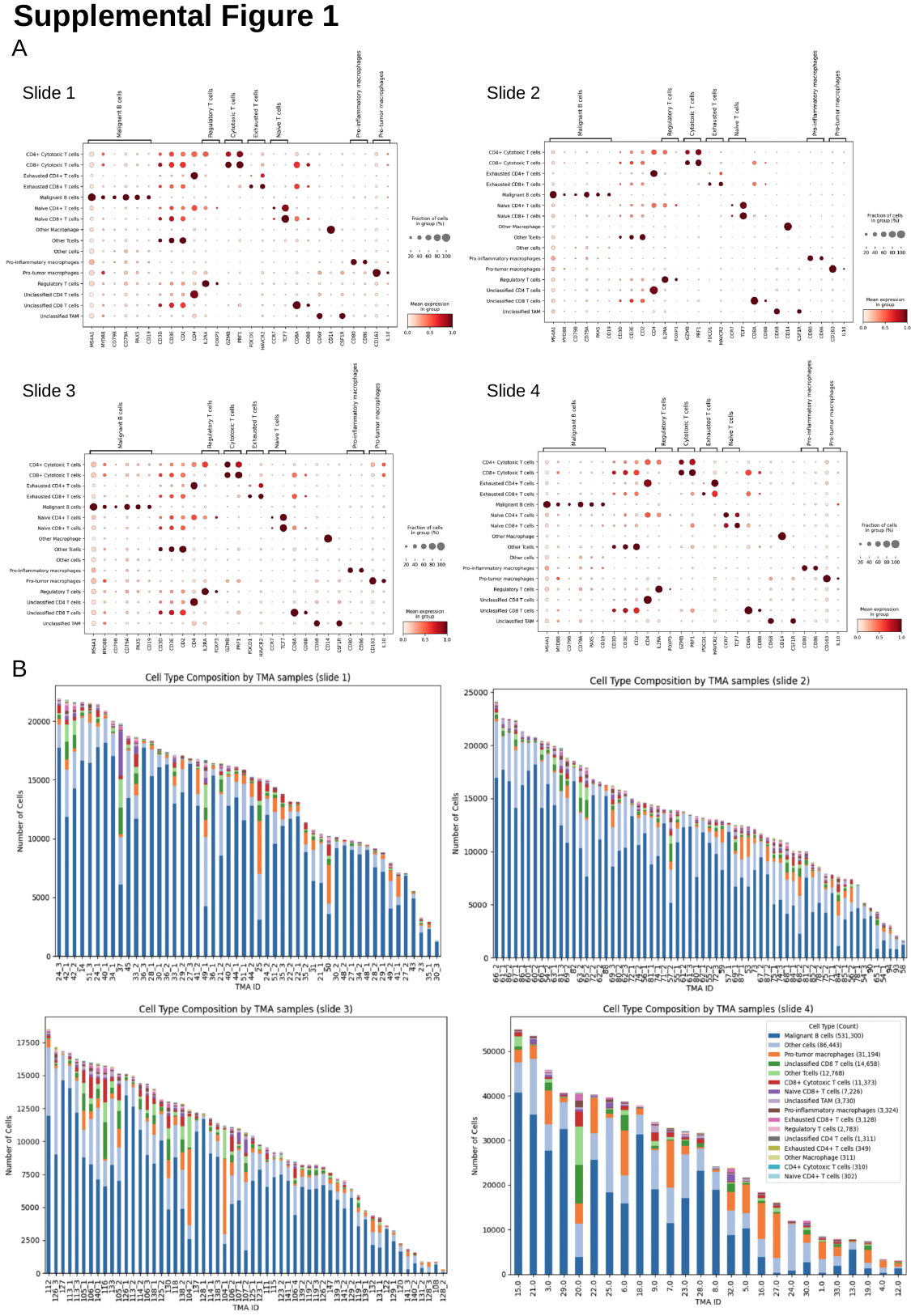

Supplemental Figure 1
A
Slide 1
Slide 2
Slide 3
Slide 4
B

### Supplementary FigureS2

#### Slide 1
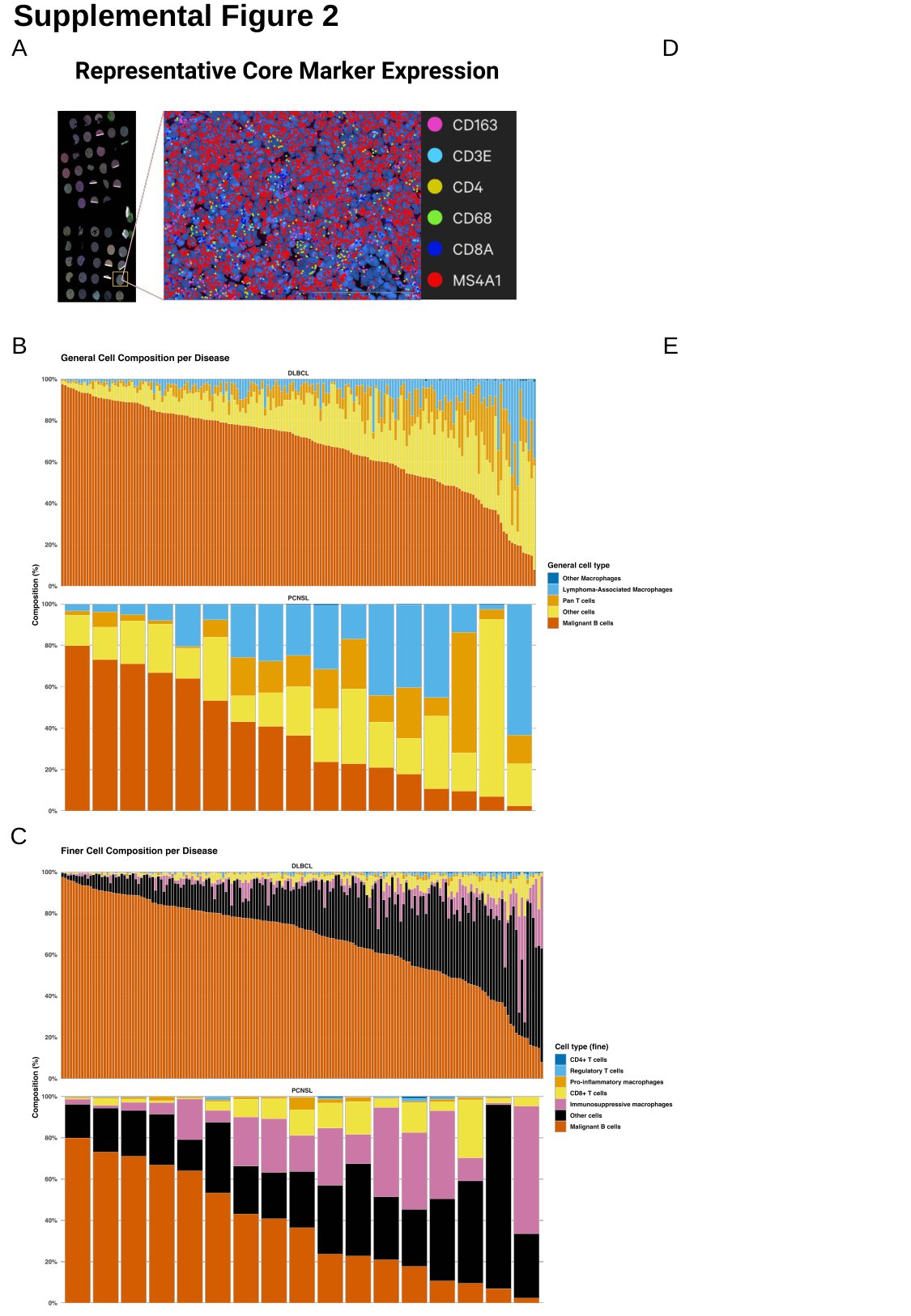

Supplemental Figure 2
D
A
B
E
C

### Supplementary FigureS3

#### Slide 1
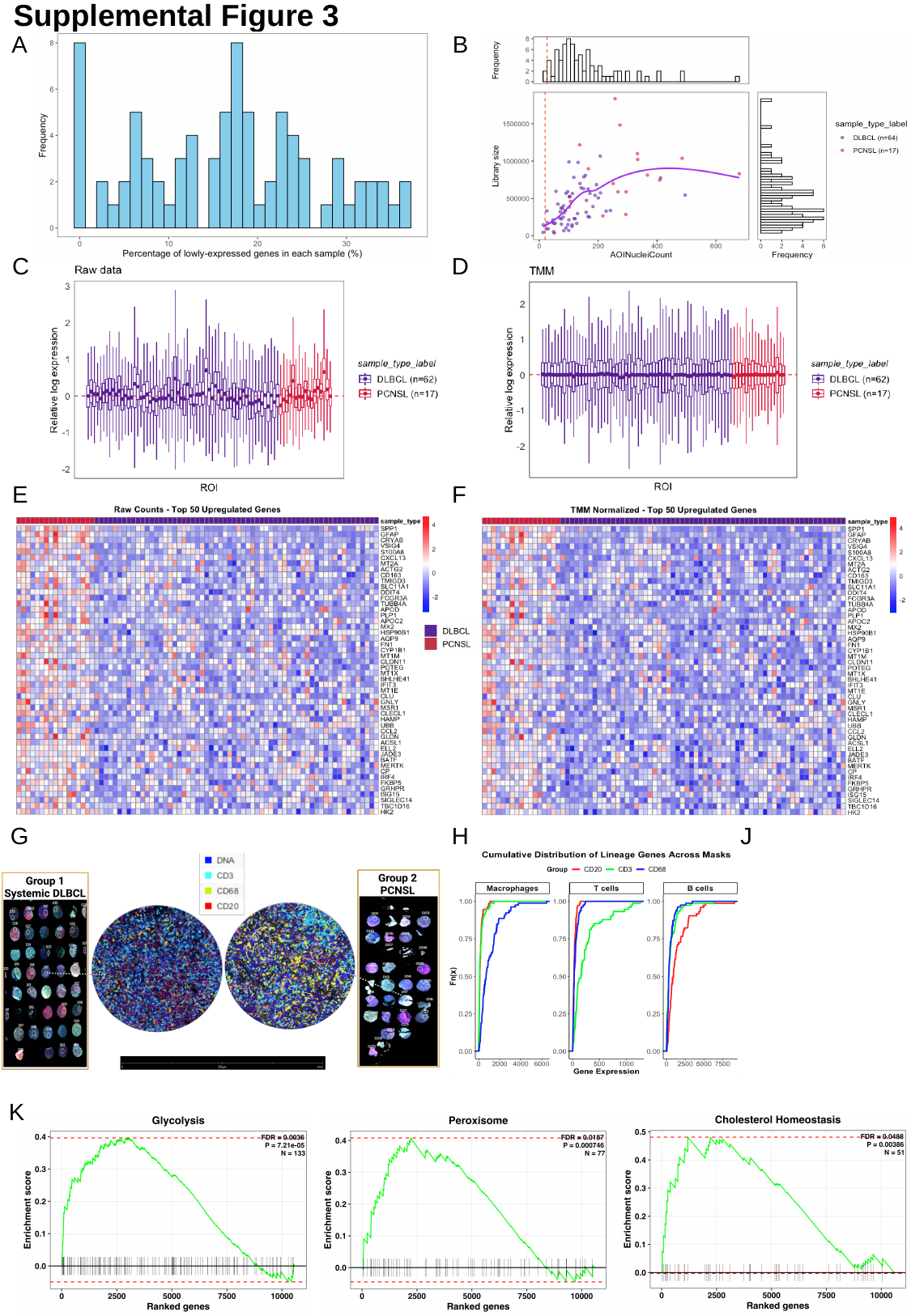

Supplemental Figure 3
B
A
C
D
E
F
H
G
H
J
K

### Supplementary FigureS4

#### Slide 1
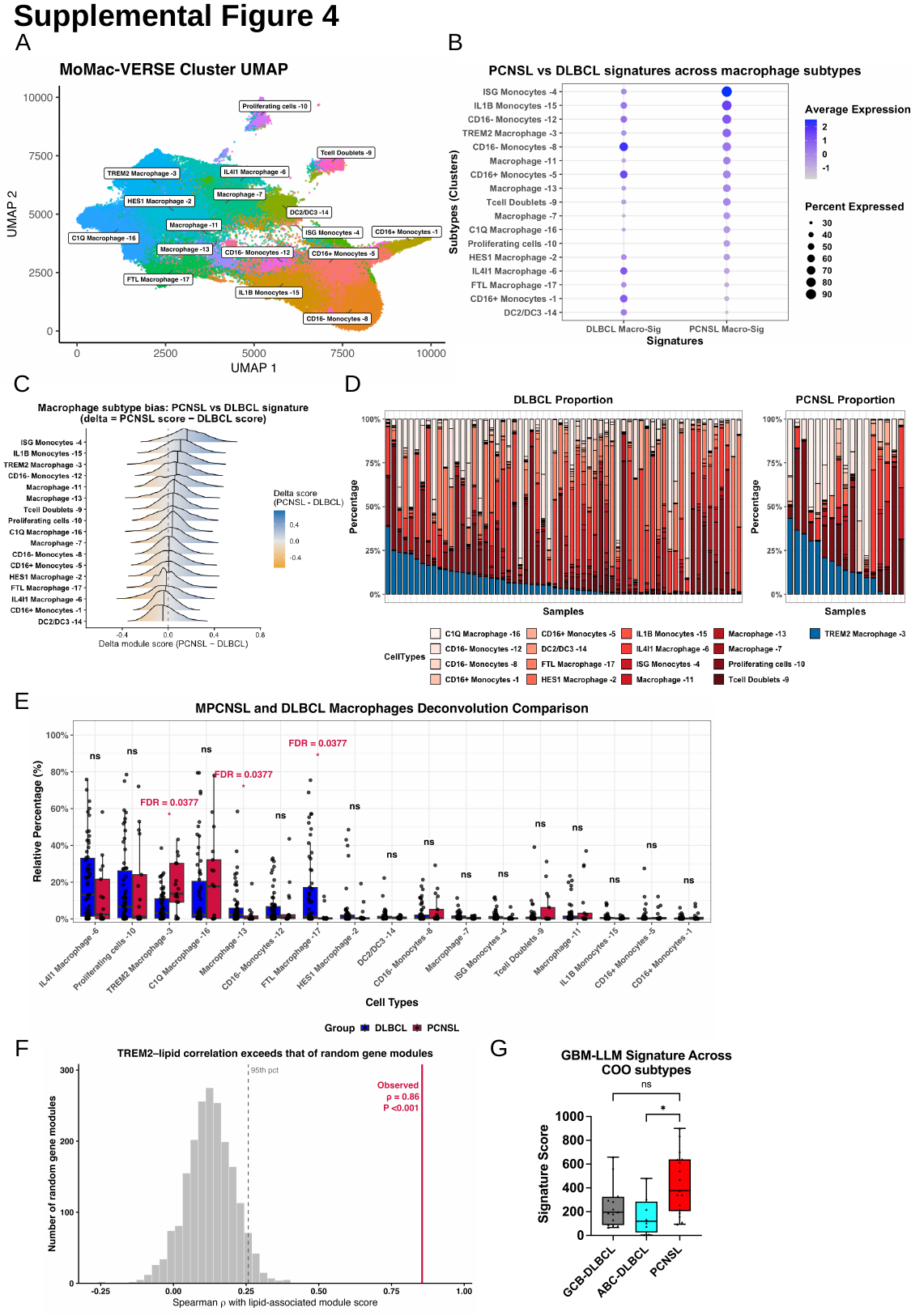

Supplemental Figure 4
A
B
C
D
E
G
F

### Supplementary FigureS5

#### Slide 1
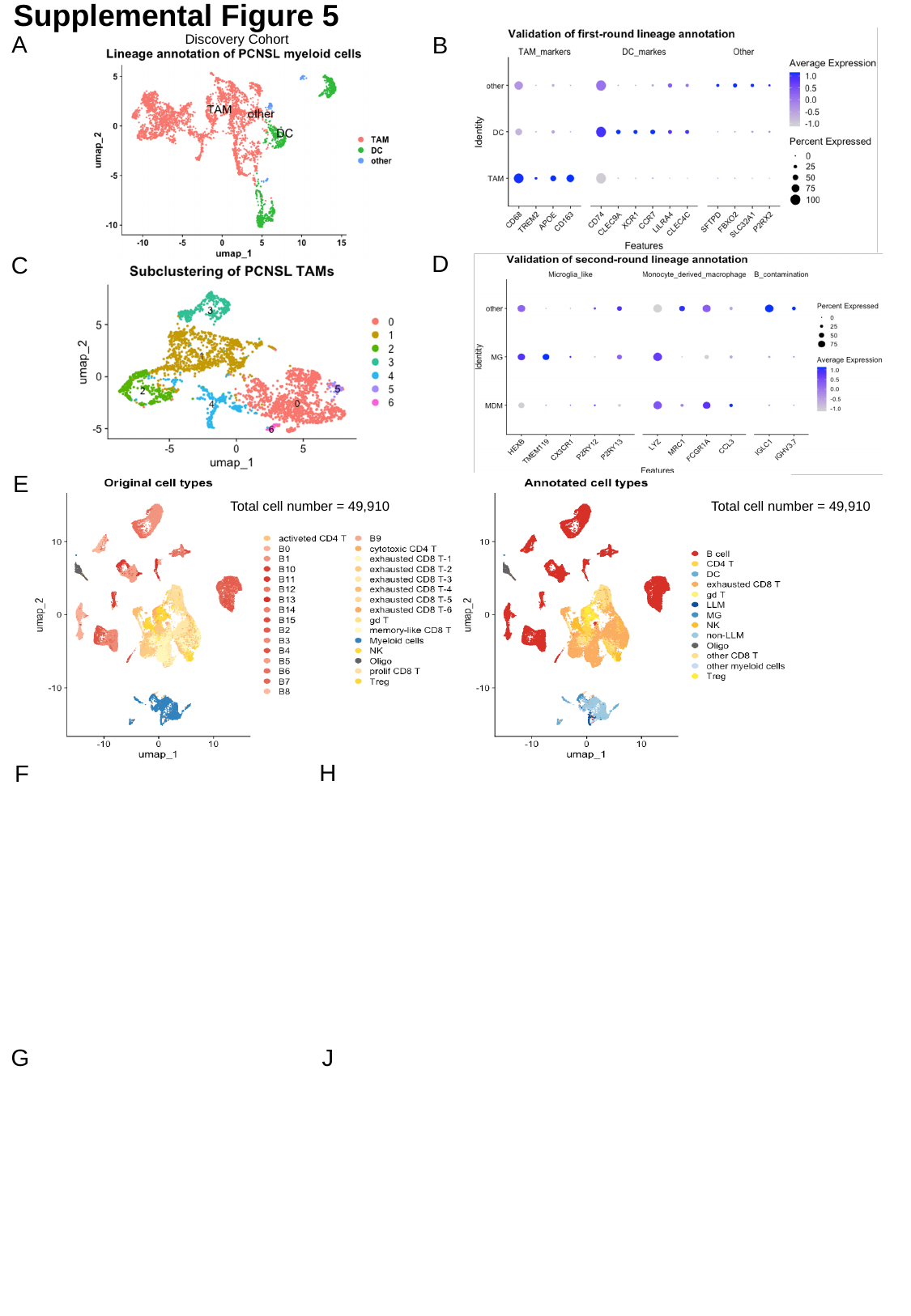

Supplemental Figure 5
A
B
Discovery Cohort
D
C
E
Total cell number = 49,910
Total cell number = 49,910
H
F
G
J

### Supplementary FigureS6

#### Slide 1
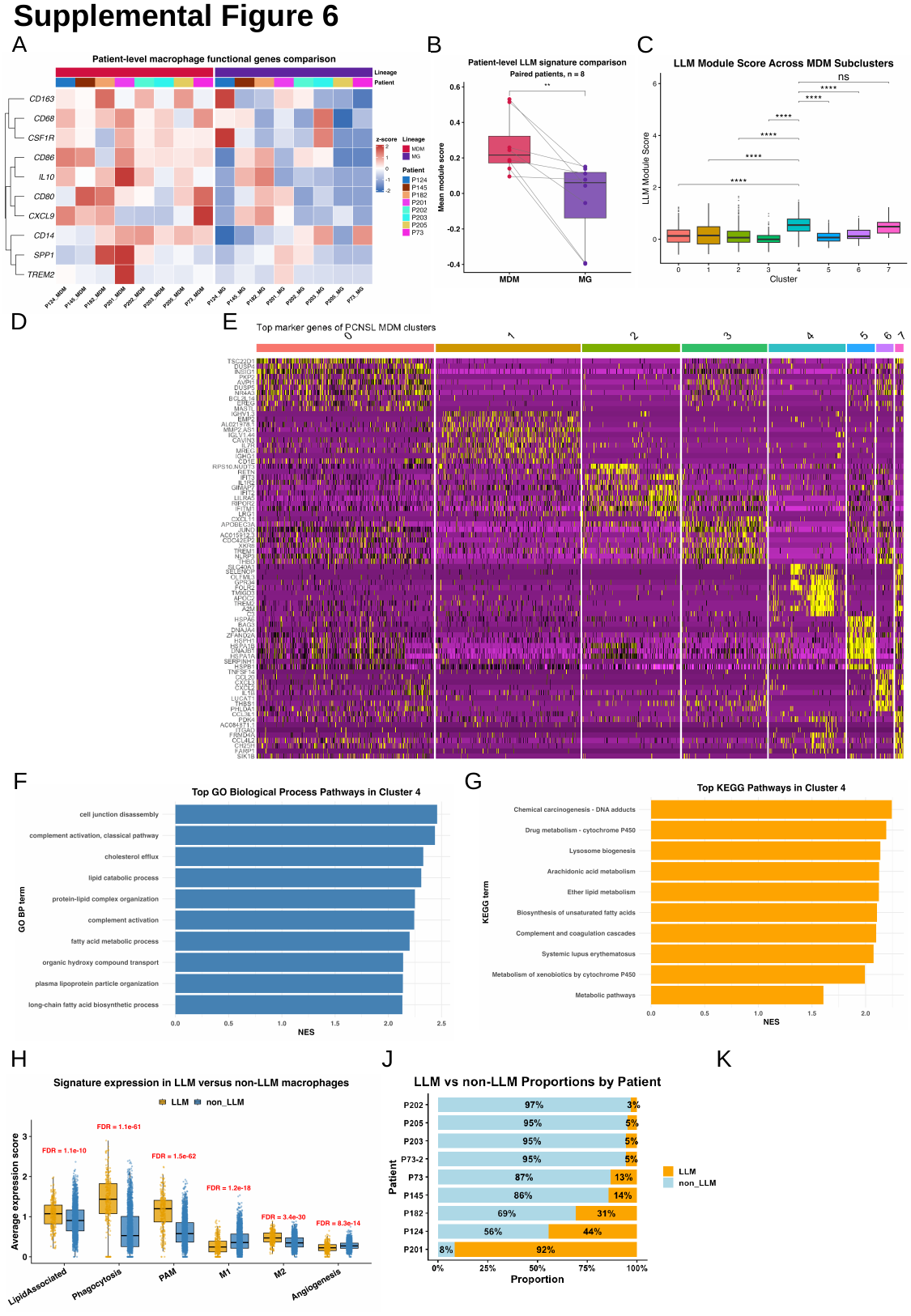

Supplemental Figure 6
B
C
A
E
D
F
F
G
H
K
J

### Supplementary FigureS7

#### Slide 1
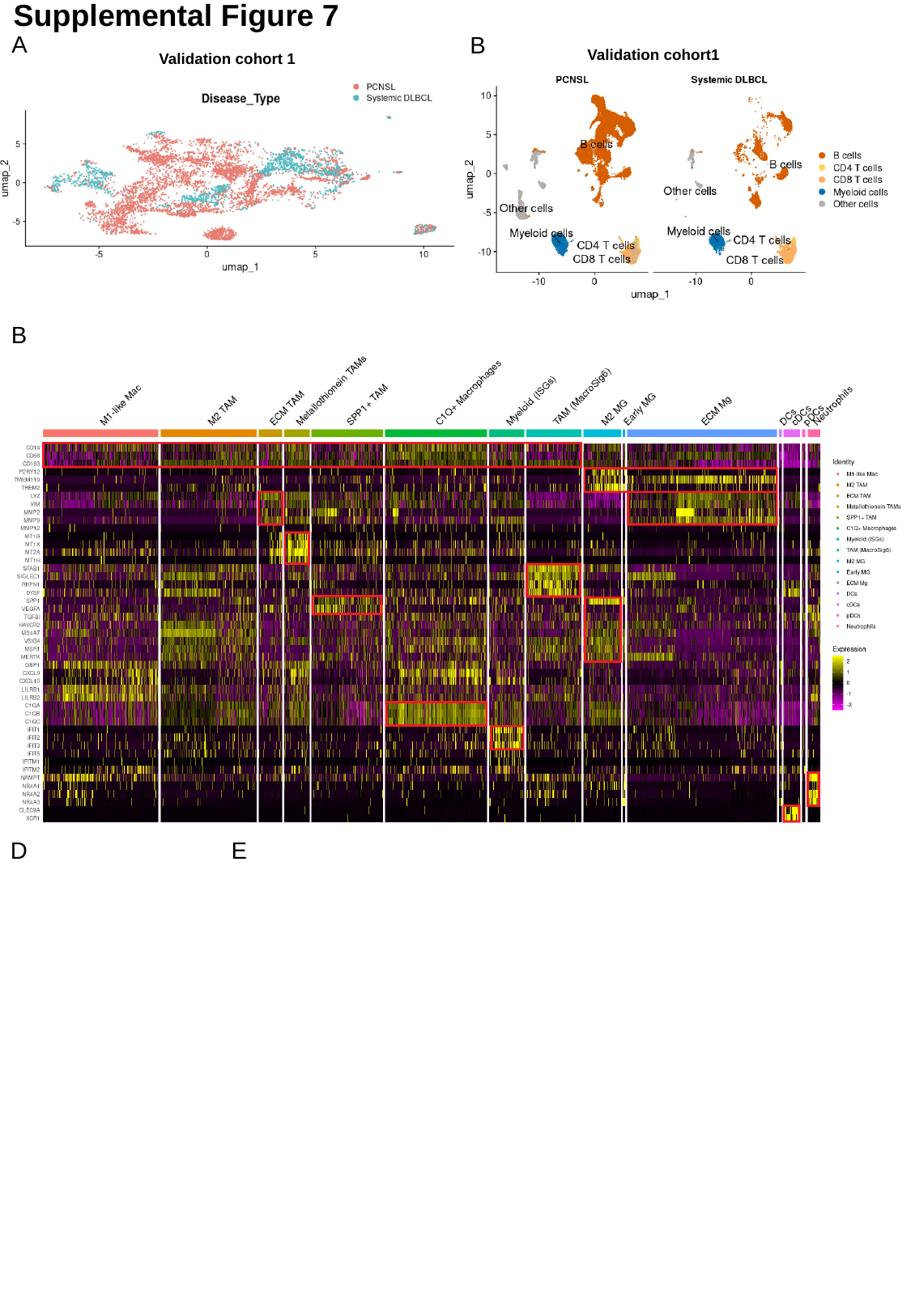

Supplemental Figure 7
A
B
Validation cohort1
Validation cohort 1
B
D
E

### Supplementary FigureS8

#### Slide 1
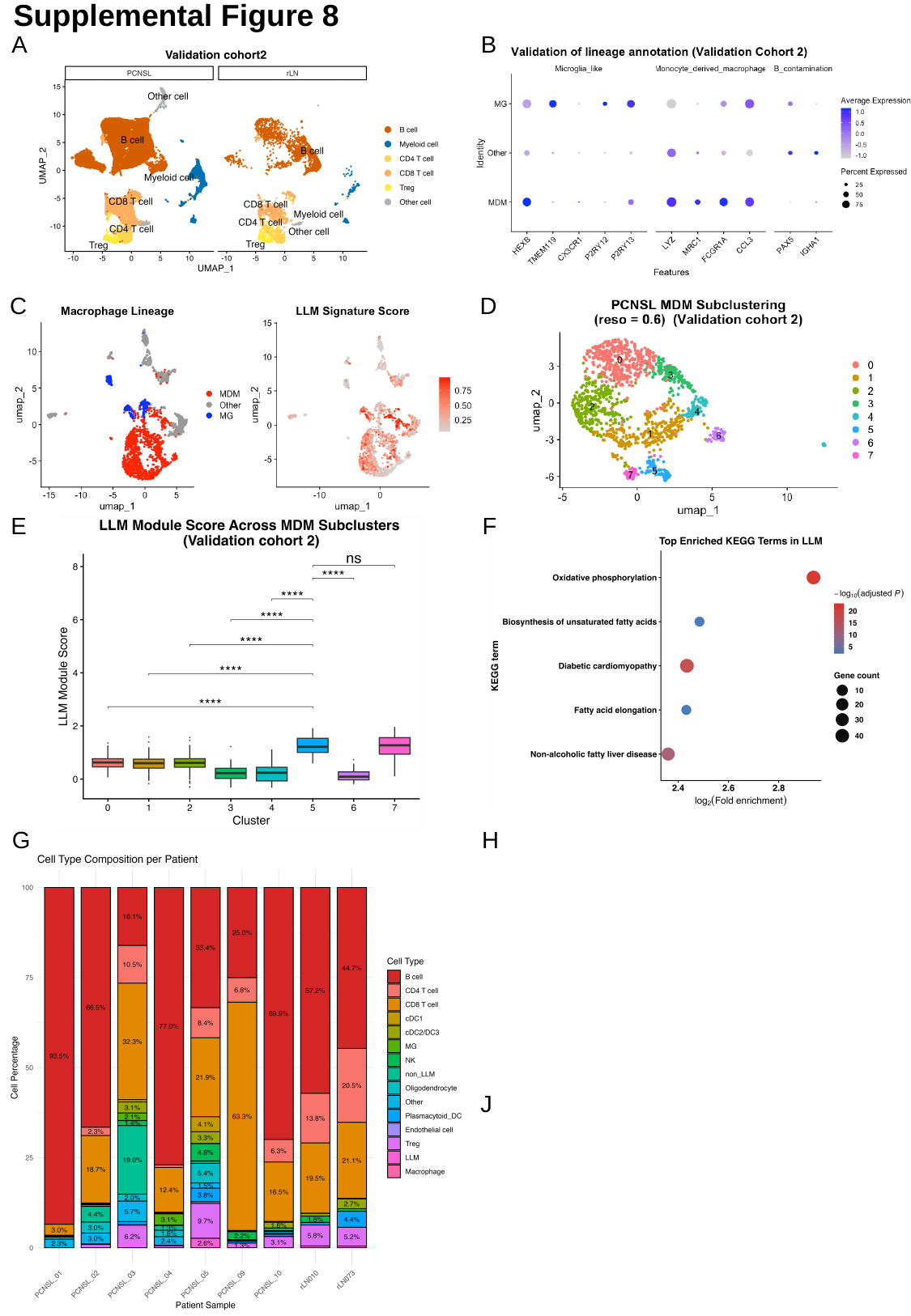

Supplemental Figure 8
B
A
C
D
E
F
G
H
J

### Supplementary FigureS9

#### Slide 1
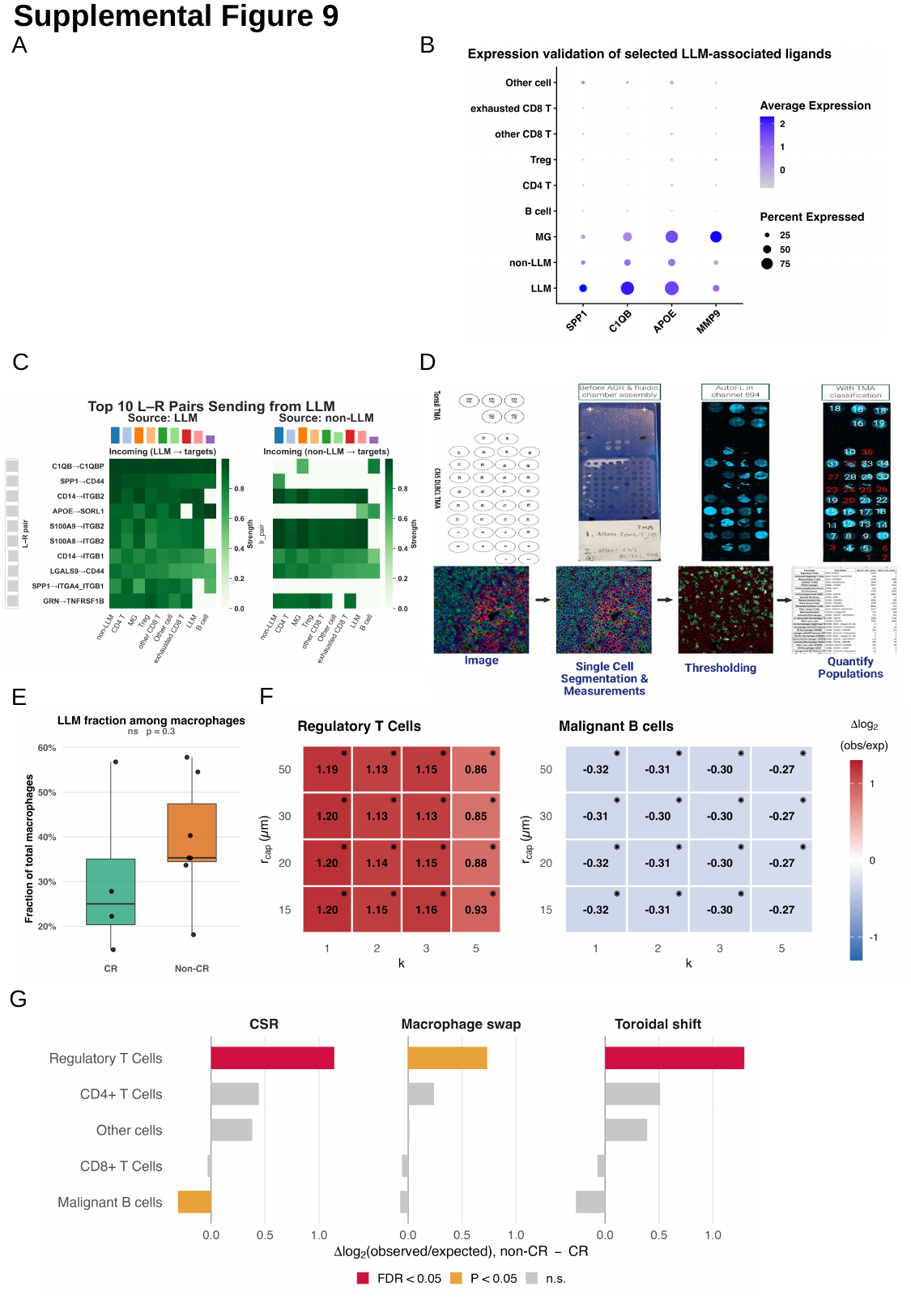

Supplemental Figure 9
A
B
C
D
F
E
G

### Supplementary FigureS10

##### Supplemental Figure 10

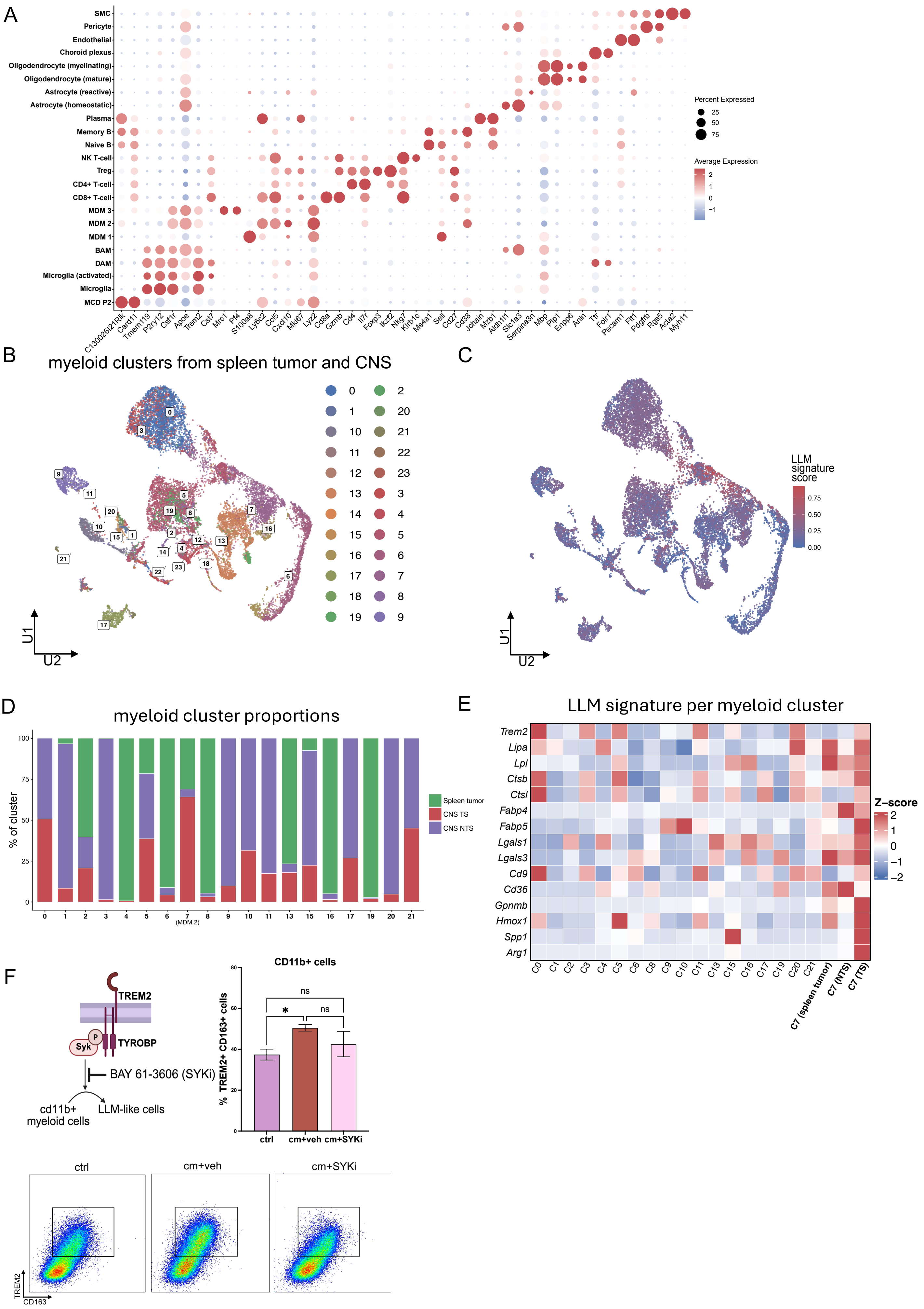
