## Supplementary Method for "A lipid-laden macrophage niche drives immunosuppression in primary central nervous system lymphoma"

*^18^Advanced Pathology Laboratory, IFOM ETS - The AIRC Institute of Molecular Oncology, Milan, Italy. Department of Oncology and Hemato-Oncology, University of Milan, Milan, Italy*

**# first authors**

***Corresponding authors:**

**Dr. Anand Devaprasath Jeyasekharan**

**Cancer Science Institute of Singapore, 14 Medical Dr, Centre for Translational**

**Medicine (MD6), Singapore 117599**

****

**Dr. Claudio Tripodo**

**Tumor Immunology Unit, Department of Sciences for Health Promotion and Mother-Child Care “G. D’Alessandro”, University of Palermo, Palermo, Italy**

**Histopathology Unit, Institute of Molecular Oncology Foundation (IFOM) ETS - The AIRC Institute of Molecular Oncology, Milan, Italy**

**Dr. Leandro Cerchietti**

**NYU Perlmutter Cancer Center**

**Division of Hematology and Oncology, Department of Medicine**

**One Park Avenue, M-036, New York, NY 10016, USA**

**SUPPLEMENTAL METHODS**

**Cell clustering and annotation of Xenium data**

Xenium cell-by-gene count matrices were imported into Seurat and subjected to standard quality control based on transcript counts, detected genes per cell and spatial distribution. Low-quality or empty cells were removed according to dataset-specific thresholds. Data were normalized, variable features were identified, and dimensionality reduction was performed using principal component analysis followed by UMAP visualization. Cells were clustered using a shared-nearest-neighbor graph-based approach, with clustering resolution selected according to the biological granularity required for downstream analysis.

Cell-type annotation was performed by integrating unsupervised clustering with canonical marker-based interpretation, as described in the main text. Cluster identities were assigned based on the expression of established lineage and cell-state markers, together with cluster-level expression patterns and spatial localization. Clusters with ambiguous marker profiles or inconsistent spatial distributions were manually reviewed and, where appropriate, merged, relabeled or retained as unresolved populations. Final cell-type labels were stored as cell-level metadata and visualized in both UMAP and tissue coordinates.

**GeoMx DSP Whole Transcriptome Atlas profiling**

Spatial transcriptomic profiling of PCNSL and DLBCL FFPE tissues was performed using the GeoMx Human NGS Whole Transcriptome Atlas (WTA) assay (GMX-RNA-NGSHuWTA-4; NanoString) according to the manufacturer’s instructions.

Freshly cut 5-µm FFPE sections were mounted on Bond Plus slides (S21.2113.A; Leica Biosystems), baked at 60°C for 1 hour, and processed using the Bond Max automated staining system. Processing included deparaffinization, rehydration, antigen retrieval with ER2 solution (AR9640; Leica Biosystems) at 100°C for 20 minutes, Proteinase K digestion (1 µg/mL for 15 minutes), and post-fixation in 10% neutral-buffered formalin for 5 minutes, followed by two 5-minute washes in formalin stop buffer.

Overnight in situ hybridization was performed using the GeoMx Human NGS WTA probe set, targeting more than 18,000 protein-coding genes. Slides were subsequently washed twice in equal volumes of 4× SSC and 100% formamide at 37°C for 25 minutes to remove nonspecific probes. After blocking with Buffer W (200 µL per slide; GMX-PREP-RNAFFPE-12; NanoString) for 30 minutes at room temperature, sections were stained with CD68-AF594 (sc-20060; Santa Cruz Biotechnology), CD3 (A0452; Dako), CD20-AF647 (NBP2-47840; Novus Biologicals), and the nuclear stain SYTO 13.

Slides were imaged using the GeoMx DSP instrument with GeoMx software v2.4.0.421. Regions of interest (ROIs) were selected under expert pathologist guidance. Fluorescence-channel thresholds were optimized for each slide to generate CD68⁺, CD3⁺, and CD20⁺ morphology masks. Each ROI was segmented into macrophage-, T-cell-, and B-cell-enriched areas of illumination (AOIs) based on the corresponding masks. AOIs were exposed to ultraviolet light, and the released indexing oligonucleotides were collected into individual wells for sequencing. Gene counts from each AOI therefore represented aggregate expression from all cells captured within the corresponding morphology-defined mask.

Collected oligonucleotides were amplified using the GeoMx Seq Code Primer Plate and Master Mix (GMX-NGS-SEQ; NanoString). PCR products were pooled and purified twice using AMPure XP beads (A63880; Beckman Coulter). Library size and concentration were assessed using the Agilent High Sensitivity DNA Kit and Bioanalyzer. Libraries were sequenced on an Illumina HiSeq 3000 or NovaSeq 6000 platform using dual indexing and 2 × 27-bp paired-end reads.

**GeoMx DSP data processing**

Raw GeoMx WTA sequencing data were processed using standR(1) with limma and vissE. Gene-level quality control was performed using standR::addPerROIQC (version ‘1.10.0’) with default parameters, which flags genes falling below a log₂ CPM threshold (4.08, derived from a minimum count of 5) in at least 90% of AOIs. No genes met this criterion, consistent with prior removal of probes below the limit of quantification during data export from the GeoMx DSP platform. AOIs with 20 or fewer nuclei were subsequently excluded, removing 3 AOIs for downstream analysis. Data quality and normalization performance were assessed using relative log expression plots, heatmaps of highly variable genes, and multidimensional scaling. Expression values were normalized using the trimmed mean of M-values method.

Cell lineage signature scores were calculated using published marker sets(2). Macrophage markers included CD68, CD163, FCGR1A, and CSF1R. T-cell markers included CD3D, CD3E, UBASH3A, CD2, and TRBC2. B-cell markers included MS4A1, CD79A, CD79B, CD19, and PAX5. Cumulative expression scores were calculated within each mask and used for downstream comparisons.

**BayesPrism deconvolution of GeoMx CD68^+^ AOIs using MoMac-VERSE**

BayesPrism(3) was used to estimate macrophage-state composition in GeoMx DSP WTA CD68^+^ macrophage-enriched AOIs using the MoMac-VERSE human monocyte/macrophage atlas as the single-cell reference. For deconvolution, raw GeoMx count matrices after gene- and AOI-level quality control were used as required by BayesPrism, whereas TMM-normalized data were used for differential expression analyses. AOIs from PCNSL and DLBCL were retained after filtering for >20 nuclei per AOI. After AOI-level QC, all 17 PCNSL AOIs and 62 of 64 DLBCL AOIs were retained for BayesPrism deconvolution.

The MoMac-VERSE RNA count matrix was extracted from the Seurat object, with MoMac cluster annotations used as both cell-type and cell-state labels. Reference genes were filtered using BayesPrism cleanup procedures to remove ribosomal, mitochondrial, MALAT1, and sex chromosome-associated genes, and genes were required to be expressed in at least 5 cells. Concordance between GeoMx bulk profiles and the single-cell reference was assessed using BayesPrism bulk-versus-single-cell diagnostic plots, and cell-type correlation and outlier-gene diagnostics were inspected before model fitting. BayesPrism objects were constructed using count matrices with common gene features between GeoMx AOIs and the MoMac-VERSE reference, with outlier.cut = 0.01 and outlier.fraction = 0.1. Final deconvolution estimates were extracted from the posterior cell-state fraction matrix (‘which.theta = "final"‘), and proportions were matched back to GeoMx AOI annotations. For focused macrophage-state comparisons, macrophage-related MoMac-VERSE clusters were retained if their overall median estimated abundance across AOIs exceeded 1%. Group comparisons were performed at the AOI level using Wilcoxon rank-sum tests with Benjamini-Hochberg correction.

**TREM2 and lipid-associated module scoring in CD68⁺ AOIs**

Module scores were computed from TMM-normalized log₂ counts per million (CPM) values of CD68⁺ AOIs, generated with edgeR (cpm(log = TRUE, prior.count = 1)).

The lipid-associated macrophage module comprised 11 genes previously reported to define lipid-handling macrophage states (TREM2, LIPA, LPL, CTSB, CTSL, FABP4, FABP5, LGALS1, LGALS3, CD9, and CD36)(4); all 11 were represented in the WTA panel.

The TREM2 macrophage module was derived from the MoMac-VERSE reference atlas(5). Cluster-specific markers were identified using presto::wilcoxauc on log-normalized expression, and genes assigned to the "TREM2 Macrophage-3" cluster were retained if they met the following criteria: Benjamini-Hochberg adjusted P < .05, area under the receiver operating characteristic curve (AUC) > 0.65, detection in at least 10% of cluster cells, and a difference in detection rate between cluster and non-cluster cells of at least 15%. Genes were ranked by AUC and the top 100 were selected, of which 85 were represented in the WTA panel. To avoid circularity, the 7 genes shared with the lipid-associated module (TREM2, LIPA, CTSB, FABP5, LGALS1, LGALS3, and CD9) were removed from the TREM2 module, leaving 78 genes for scoring. The full gene list is provided in supplemental Table1.

For each module, expression values were standardized gene-wise across AOIs (z score), and the module score for each AOI was defined as the mean z score across module genes. Scores were therefore centered at 0, with positive values indicating above-average expression of the module relative to the other AOIs in the cohort.

Association between the 2 module scores was assessed across the 17 PCNSL CD68⁺ AOIs (1 AOI per patient) using the Spearman rank correlation. To test whether the observed correlation was specific to the TREM2 gene set rather than a general property of any macrophage-associated gene module, we performed a permutation analysis: 2000 random gene sets of the same size (n = 78) were sampled without replacement from all genes in the WTA panel, excluding the 11 lipid-associated module genes. Each random set was scored identically and correlated with the lipid-associated module score. The empirical P value was calculated as (number of random sets with |ρ| ≥ |ρ_observed| + 1) / (2000 + 1). Analyses were performed in R version 4.3 with a fixed random seed (42) to ensure reproducibility.

**Single-cell RNA-seq sample preparation**

Tissue samples of the University of Queensland single-cell cohort were obtained from Plymouth NHS Trust as part of BRAIN UK, which is supported by Brain Tumour Research and has been established with the support of the British Neuropathological Society and the Medical Research Council(6). Single-cell RNA-seq was performed on 13 FFPE PCNSL biopsies using the 10x Genomics Flex Gene Expression 16-plex assay. One 25 µm scroll per sample was deparaffinized in xylene and dissociated according to the manufacturer’s FFPE tissue protocol using a gentleMACS Octo Dissociator. Nuclei concentration was determined using AOPI staining before probe hybridization. After 20 hours of hybridization, samples underwent individual washing and pooling. Nuclei were loaded onto the Chromium X platform with a target of 8,000 cells per sample. Libraries were sequenced by the Australian Genome Research Facility on an Illumina NovaSeq X platform, targeting 30,000 reads per nucleus.

**Single-cell RNA-seq preprocessing and annotation**

For published datasets, initial filtering and quality-control procedures followed the corresponding studies. For the PCNSL discovery dataset, the Seurat object was processed using a Seurat-based workflow. Gene expression was log-normalized using ‘NormalizeData‘ with a scale factor of 10,000, and the top 2,000 variable features were identified using the vst method. Data were scaled using variable features, followed by principal component analysis, nearest-neighbor graph construction, UMAP visualization, and Louvain clustering. Unless otherwise specified, dimensionality reduction and clustering were performed using the first 40 principal components.

Major cell populations were annotated using author-provided labels where available, cluster-defining genes identified by ‘FindAllMarkers‘, and canonical lineage markers. The myeloid compartment was subset and reprocessed independently to resolve tumor-associated macrophages, dendritic cells, and other myeloid populations. Tumor-associated macrophages were further subset and reclustered to distinguish monocyte-derived macrophages (MDMs) from microglia-like cells using marker programs including ‘LYZ‘, ‘MRC1‘, ‘FCGR1A‘, and ‘CCL3‘ for MDMs and ‘HEXB‘, ‘TMEM119‘, ‘CX3CR1‘, ‘P2RY12‘, and ‘P2RY13‘ for microglia-like cells. Clusters with immunoglobulin gene enrichment were annotated as likely B-cell contamination or other cells and excluded from macrophage-focused analyses where appropriate.

The lipid-associated module captures a canonical lipid-handling program described largely in chronic inflammatory and metabolic settings, whereas the GBM-derived signature reflects a lipid-laden macrophage state arising specifically within CNS tumors. Because these gene sets capture complementary aspects of the same phenotype, and because TREM2, a defining LAM marker, was itself enriched in PCNSL, we combined them into a single LLM signature for the single-cell analyses.

LLM signature scores were calculated using Seurat ‘AddModuleScore‘ based on the union of lipid-associated macrophage genes (‘TREM2‘, ‘LIPA‘, ‘LPL‘, ‘CTSB‘, ‘CTSL‘, ‘FABP4‘, ‘FABP5‘, ‘LGALS1‘, ‘LGALS3‘, ‘CD9‘, ‘CD36‘)(7) and GBM-associated LLM genes (‘GPNMB‘, ‘FABP5‘, ‘HMOX1‘, ‘SPP1‘, ‘ARG1‘)(8). After removing the gene shared between the two sets (FABP5), 15 unique genes were used; only genes detected in each dataset were retained for scoring. Patient-level comparisons were performed by averaging cell-level module scores within each patient and annotated lineage. PCNSL MDMs were reclustered, and the cluster with the highest LLM signature activity was annotated as the LLM cluster. Differential markers of LLMs versus non-LLM macrophages were identified using Seurat ‘FindMarkers‘ with Wilcoxon testing, ‘min.pct = 0.25‘, and ‘logfc.threshold = 0.25‘.

For pathway analysis, LLM-upregulated genes were defined using adjusted P < 0.05 and average log2 fold change > 0.58. Gene identifiers were converted from symbols to Entrez IDs using ‘org.Hs.eg.db‘. Over-representation analysis was performed using ‘clusterProfiler‘ for GO Biological Process and KEGG pathways and ‘ReactomePA‘ for Reactome pathways, using the tested macrophage gene set as the background universe where applicable. Ranked GSEA was performed using log2 fold change-ranked genes with Benjamini-Hochberg correction.

**LIANA+ ligand-receptor analysis**

Cell-cell communication analysis was performed using LIANA+(9) on the discovery PCNSL single-cell RNA-seq cohort. The processed AnnData object was used as input, and log-normalized expression values from the 'logcounts' layer were assigned as the expression matrix for ligand-receptor inference. Cell populations were grouped using the curated 'celltype_cellchat' annotation. The LIANA+ consensus ligand-receptor resource was used, and ligand-receptor interactions were retained only when the ligand and receptor were expressed in at least 10% of cells in the corresponding source or target population ('expr_prop = 0.10').

Two complementary LIANA+ methods were applied. First, CellPhoneDB inference was performed to estimate ligand-receptor mean expression and permutation-based interaction significance. CellPhoneDB-derived plots were filtered using 'cellphone_pvals <= 0.05'. Second, LIANA+ rank aggregation was used to integrate ligand-receptor evidence across methods and generate consensus interaction rankings. Rank-aggregate results were exported for downstream visualization and comparison.

The analysis focused on LLM and non-LLM macrophages as source populations. Target populations included B cells, CD4^+^ T cells, Tregs, exhausted CD8^+^ T cells, other CD8^+^ T cells, other cells, LLMs, non-LLM macrophages, and microglia. For overview visualizations, the top ligand-receptor interactions were selected by magnitude rank, with lower magnitude ranks indicating stronger inferred interactions. Interaction specificity was assessed using LIANA+ specificity rank, and visualizations were filtered using specificity-rank thresholds as indicated.

To compare LLM-derived and non-LLM-derived communication programs, we identified ligand-receptor-target combinations detected in both LLM and non-LLM source populations. Combinations were retained if either source population showed a specificity rank below 0.20. Retained combinations were ranked by the absolute difference in magnitude rank between the 2 source populations, and the top 10 were visualized as differential dot plots. Combinations inferred in only one source population were not included, as magnitude ranks could not be compared directly.

For heatmap-based summaries, an interaction strength score was calculated as (1 − magnitude rank) × (1 − specificity rank), such that stronger and more specific interactions received higher values. Outgoing matrices summarized interaction strength by source population, and incoming matrices summarized interaction patterns by target population. For LLM-focused analyses, the top 10 LLM-derived ligand-receptor pairs were selected by summed interaction strength across target populations and compared with the corresponding non-LLM-derived interactions. Heatmaps were generated without row or column scaling, preserving relative inferred interaction strength across ligand-receptor pairs and target populations.

**CellScape tissue preparation and imaging**

Multiplex immunofluorescence staining and imaging were performed on the CellScape Precise Spatial Proteomics platform (Bruker). FFPE sections were mounted on standard histology slides and deparaffinized according to the study protocol. Antigen retrieval was performed in CC1 buffer (Roche, 950-124) at 95°C for 20 minutes using a pressure-based retrieval system. Flow chambers were assembled using the CellScape Slide Assembly Tool (Bruker, MAN-10200-01) to mount Whole Slide Imaging Chamber coverslips (Bruker, PRSM-CS-WSIC-010).

Before staining, a baseline autofluorescence scan was acquired to assess tissue integrity. Regions with tissue folding or detachment were excluded. Antibodies were diluted in a storage buffer (Bruker, PRSM-BUF-STR-50 mL). After each staining and imaging cycle, tissues were washed, photobleached using EpicIF solution (Bruker, PRSM-BUF-EPIC-250 mL), and washed again. A background scan was acquired before each cycle. Image stacks were aligned using CellScape Navigator software. The output consisted of 16-bit OME-TIFF images with automated stitching, alignment, and flat-field correction. Pixel-wise autofluorescence correction was applied independently for each marker using cycle-specific autofluorescence images.

**CellScape reinterrogation cycles**

After completion of the initial antibody panel, slides were stored at 4°C in a storage buffer for two months. Reinterrogation imaging was performed on the same instrument using additional markers under identical wash, photobleach, and acquisition conditions. Unlabeled antibodies targeting TREM2 and CD36 were labeled using FluoTag-X2 smart secondaries from nanoTag Biotechnologies according to the manufacturer’s protocol.

**Cell segmentation and marker calling**

Image visualization and cell segmentation were performed in QuPath v0.5. Tissue microarray cores were manually annotated. Segmentation was performed using the Cellpose cyto3 model through the QuPath Cellpose extension. The membrane input channel was generated as a maximum projection of ATP1A1, B2M, CD45, and SMA. Nuclear detection was performed using the Sytox Orange channel. For each detected cell, cell area, centroid coordinates, and mean marker intensities were exported for downstream analysis in Python and R.

Marker positivity was determined independently within each tissue microarray core using cell-level mean fluorescence intensities. Otsu thresholding was used for most markers. For highly abundant markers, including CD20 and CD45, Otsu thresholding was applied after log transformation of intensity values. As macrophages and T-cells are both immune-lineage populations and may be closely apposed within dense DLBCL tissue, lineage-specific markers such as CD68 and/or CD163 for macrophages and CD3 and/or CD8 for T-cells will be used primarily for downstream cell-type annotation in spatial transcriptomics. To minimize transcript spillover and incorrect cell assignment, we will apply conservative boundary quality control, doublet filtering, exclusion or flagging of ambiguous cell masks, and post-segmentation validation using canonical lineage transcripts and protein markers where applicable.

Cells positive for markers from more than one lineage were evaluated using within-core marker intensity quantile ranks. For each multi-positive cell, the marker with the highest quantile rank was considered the candidate true lineage marker, whereas lower-ranking markers were treated as potential spillover-derived false positives. Cells with unresolved multi-lineage positivity after correction were classified as other and were excluded from lineage-focused analyses.

**CellScape marker selection and spatial analysis**

The custom CellScape antibody panel included CD68 and the lipid-associated macrophage markers GPNMB, TREM2, and CD36. During image review and threshold optimization, TREM2 and CD36 showed higher background, greater inter-core variability, and less consistent signal-to-noise performance in FFPE PCNSL TMA cores. GPNMB showed the most reproducible macrophage-associated staining with lower nonspecific background. GPNMB⁺CD68⁺ macrophages were therefore used as a conservative protein-level proxy for LLM-like macrophages in downstream spatial analyses.

LLM-centered spatial organization was assessed using cross-type pair-correlation functions and nearest-neighbor enrichment metrics based on per-cell coordinates and phenotype annotations. LLM-like macrophages were used as the reference population, and CD4⁺ T cells, CD8⁺ T cells, regulatory T cells, and malignant B cells were evaluated as query populations. Pair-correlation functions were calculated in spatstat (v3.6-1) at 5-µm intervals up to 50 µm using translation edge correction and random-relabeling simulations. ROIs were required to contain at least 20 LLM-like macrophages, 20 CD4⁺ T cells, and 20 CD8⁺ T cells.

Nearest-neighbor enrichment analysis identified the two nearest non-LLM cells within 30 µm of each LLM-like macrophage. Boundary guarding and density-based ROI filtering were applied, and observed neighborhood frequencies were compared with simulation-based complete spatial randomness null models. Enrichment was summarized using observed and expected rates, z scores, log₂ fold changes, and empirical P values.

For treatment-response analyses, spatial metrics were initially calculated at the ROI/core level. Measurements from multiple ROIs or sections belonging to the same patient were averaged to generate one patient-level estimate. Complete response and non-complete response groups were compared using two-sided Wilcoxon rank-sum tests, with Benjamini-Hochberg correction across spatial features.

***In-vivo* murine MCD-DLBCL model**

DLBCL cells harboring MYD88^L252P^ and CD79B mutations with BCL2 overexpression (MCD_P2) were cultured in RPMI supplemented with 20% fetal bovine serum (FBS; Gibco, A4766801), 1% penicillin-streptomycin (Gibco, 15140122), 1% non-essential amino acids (Quality Biological, 116-078-721), 1% sodium pyruvate (Corning, 25-000-CI), 1% GlutaMAX (Gibco, 35050061), and 1% HEPES buffer (Gibco, 15630080) at 37°C and 5% CO₂.

For modeling PCNSL, MCD_P2 cells were resuspended in PBS at 50,000 cells/µl (>90% viability by trypan blue exclusion) and 100,000 cells were implanted via intracranial injection unilaterally into the brain parenchyma of C57BL/6 wild-type mice (12 months old males). Mice were anesthetized with 2% isoflurane and placed on a heating pad to maintain body temperature. Stereotaxic coordinates for all injections were: AP +0.4 mm, ML +2.0 mm, DV −2.5 mm and −2.4 mm. Cells were injected at 1 µl/min at each DV coordinate using a 26s-gauge needle and 5 µl Hamilton syringe on a Kopf stereotaxic frame. The needle was held in place for 5 min and withdrawn at 0.5-1.0 mm/min.

For modeling systemic DLBCL, our previously established splenic murine lymphoma model was used (Zamponi et al., manuscript under review). Briefly, C57BL/6 wild-type mice (3-4 months old males) were anesthetized with 2% isoflurane and placed on a heating pad. A subcostal incision (~1.5 cm) was made on the left flank to expose the spleen. MCD_P2 cells (1 × 10⁶ viable cells in 100 µl PBS; >90% viability by trypan blue exclusion) were injected into the distal pole of the spleen using a 27G needle and Hamilton syringe. Wet gauze was applied during needle withdrawal, followed by gentle compression for 1 minute to prevent leakage. The peritoneum was closed with absorbable sutures, and the skin with wound clips. Postoperative analgesia consisted of topical bupivacaine (0.12%) and subcutaneous meloxicam (2 mg/kg) and buprenorphine (0.5 mg/kg) at the time of surgery, followed by meloxicam (2 mg/kg) daily for 72 hours. Mice were monitored daily for weight loss and clinical signs of disease progression. Tumor burden was assessed every other day by bioluminescence imaging using an IVIS Spectrum system (PerkinElmer). Mice were anesthetized with isoflurane and injected intraperitoneally with D-luciferin (100 μL at 15 mg/mL; Revvity, 122799A), and images were acquired 10 minutes after injection with a 1 min exposure time. Bioluminescent signal was quantified using Living Image software (PerkinElmer).

**Mice tumor dissection**

MCD_P2 implanted mice were euthanized 15 days post implantation by CO₂ exposure and brains were rapidly dissected. A 4-5 mm coronal section encompassing the tumor implantation site (~2 mm on either side) was cut using a sterile scalpel and divided along the midline into the implanted tumor side (TS) and the contralateral non-tumor side (NTS). Each region was mechanically minced using sterile scalpels and enzymatically dissociated using the Neural Tissue Dissociation Kit (P) (Miltenyi Biotec, 130-092-628) per the manufacturer's protocol. The resulting suspension was passed through a 40 µm cell strainer and centrifuged at 350 × g for 5 min at 4°C. Myelin and cellular debris were removed using Debris Removal Solution (Miltenyi Biotec, 130-109-398), and erythrocytes were lysed with RBC Lysis Buffer (Invitrogen, 00-4333-57) for 5 min at room temperature, quenched with PBS, and centrifuged at 350 × g for 5 min. Cells were passed through a 30 µm filter twice to remove any residual clumped cells. Spleens were dissected 15 days post implantation and mechanically dissociated by passage through a 70 µm cell strainer using the plunger of a 1 ml syringe. The resulting cell suspension was centrifuged at 350 × g for 5 min, and erythrocytes were lysed with RBC Lysis Buffer (Invitrogen, 00-4333-57) for 5 min at room temperature, followed by centrifugation at 350 × g for 5 min. For each model, an equal number of cells were pooled from n=3 mice before single-cell RNA sequencing.

**Murine single-cell RNA sequencing**

Libraries were generated using the 10x Genomics Single Cell 3' v4 (polyA) chemistry. Reads were aligned to the mm10-2020-A mouse reference genome using Cell Ranger (v9.0.0). Data integration and preprocessing. Single-cell RNA-seq data from CNS (non-tumor site, NTS, and tumor site, TS) and spleen samples were integrated using Seurat v5.5.0. Gene expression was normalized using a combination of standard log-normalization for most exploratory analyses and analytic Pearson residuals (Lause, Berens & Kobak, 2021, Genome Biology). Dimensionality reduction and integration. Principal component analysis was performed on Pearson residuals using irlba::prcomp_irlba (20 PCs), with residuals clipped at ±√n per the recommended approach to control outlier influence. Batch correction across tissue/dataset origin was performed with Harmony, using a combined batch variable capturing both datasets (CNS/Spleen) and tissue context (NTS/TS) to correct residual technical variation within the CNS compartment. UMAP embeddings were computed on Harmony-corrected coordinates, and Louvain clustering (FindClusters) was used to define joint myeloid clusters spanning CNS and spleen. Cell type identification. Cell types were annotated using canonical marker genes (e.g., Tmem119, P2ry12, Cx3cr1 for microglia; Apoe, Trem2 for activated myeloid; C130026I21Rik, Card11 for tumor/MCD_P2 lymphoma cells) combined with automated label transfer and module scoring (UCell). Putative monocyte-derived macrophage populations (MDM 1/MDM 2/MDM 3) were distinguished from resident microglia/DAM/BAM based on differential expression and validated against an independent orthogonal reference: the Mouse Cell Atlas (Han et al., Microwell-seq). Differential expression and pathway analysis. Pairwise differential expression was performed using the Wilcoxon rank-sum test (Seurat::FindMarkers/FindAllMarkers), with genes filtered for minimum percent expression (≥5–10%) and exclusion of mitochondrial, ribosomal, and predicted/non-coding genes (Gm prefix) prior to ranking. Gene set enrichment analysis (GSEA) was performed using fgsea against Hallmark, KEGG (legacy), and GO Biological Process gene sets (MSigDB, accessed via msigdbr), ranked by avg_log2FC for two-group comparisons. Pathway activity was additionally estimated using PROGENy (14 core signaling pathways) via decoupleR::run_wmean, and single-cell-resolution gene set scoring was performed using UCell (population-level) and AUCell-style scoring via decoupleR::run_aucell (per-cell resolution, enabling Wilcoxon-based statistical comparison of pathway activity between specific cell populations). Statistical testing. All pairwise group comparisons of UCell/AUCell scores used the two-sided Wilcoxon rank-sum test; multiple-comparison correction was performed using the Benjamini-Hochberg (BH) procedure unless otherwise noted. Software environment. All analyses were performed in R 4.6.0 using Seurat 5.5.0, CellChat 2.1.2, decoupleR 2.17.0, fgsea 1.38.0, msigdbr 26.1.0, ComplexHeatmap 2.28.0, and ggplot2 4.0.3.

**Murine primary myeloid cell and MCD_P2 co-culture system**

Adult C57BL/6 wild-type mice (3-4 months old males) were euthanized by CO₂ exposure and brains were rapidly dissected. The cerebellum and olfactory bulbs were removed, and forebrain tissue was mechanically minced, enzymatically dissociated, and cleared of myelin, cellular debris, and erythrocytes as described above. Cells were labeled with CD11b MicroBeads (Miltenyi Biotec, 130-097-142) for 15 minutes at 4°C, washed, and resuspended in MACS buffer prior to positive selection on an LS column (Miltenyi Biotec, 130-042-401). CD11b+ cells were eluted, centrifuged at 350 × g for 5 minutes, and plated at a concentration of 2.5 × 10⁵ cells/ml in Microglial Medium (ScienCell, 1901).

To generate conditioned medium, MCD_P2 cells were cultured at 1 × 10⁶ cells/ml in RPMI supplemented with 5% dialyzed FBS for 48 hours. The supernatant was collected by centrifugation at 350 × g for 5 minutes, passed through a 0.2 µm syringe filter, and applied to adherent microglia at 80% confluency for 5 days. Control microglia received RPMI supplemented with 5% dialyzed FBS alone for the same duration. Where indicated, the SYK inhibitor BAY 61-3606 dihydrochloride (SYKi; 1 µM, SelleckChem, S7006) or an equivalent volume of DMSO was added concurrently for 5 days, with media replenished every 2 days. SYKi was withdrawn prior to tumor cell addition to specifically assess the effect of microglial polarization state on tumor cell proliferation.

MCD_P2 cells were labeled with CellTrace Violet (Life Technologies, C34557) per the manufacturer's instructions and seeded onto pre-conditioned microglia at 2.5 × 10⁵ cells/ml. Co-cultures were maintained for 36 hours, after which adherent cells were detached with Accutase (Invitrogen, 00-4555-56) for 10 minutes at 37°C, pooled with non-adherent cells from the same well, and processed for flow cytometric analysis.

**Flow Cytometry**

The MDM-like transformation of CD11b^+^ primary cells treated with MCD_P2 derived conditioned media with 1μM SYKi or vehicle for 5 days was determined using CD11b (BioLegend, 101226), TREM2 (R&D Systems, FAB17291A) and CD163 (BioLegend, 155325). CellTrace Violet was used to identify MCD_P2 and assess proliferation. Flow cytometry data were acquired using a BD FACSymphony™ A1 Cell Analyzer (BD Biosciences) and analyzed using FlowJo™ v10.8.0. Experiments were performed using 3-6 independent biological replicates per condition, as indicated in the figure legends.

**SUPPLEMENTAL METHODS REFERENCES**
